# Modified Hypoxia Inducible Factor expression in CD8+ T cells increases anti-tumor efficacy

**DOI:** 10.1101/2020.06.18.159137

**Authors:** Pedro Veliça, Pedro Pacheco Cunha, Nikola Vojnovic, Iosifina Petrina Foskolou, David Bargiela, Milos Gojkovic, Helene Rundqvist, Randall Scott Johnson

## Abstract

Adoptive transfer of anti-tumor cytotoxic T cells is a novel form of cancer immunotherapy, and a key challenge is to ensure the survival and function of the transferred T cells. Immune cell survival requires adaptation to different micro-environments, and particularly to the hypoxic milieu of solid tumors. The HIF transcription factors are an essential aspect of this adaptation, and we undertook experiments to define structural determinants of HIF that would potentiate anti-tumor efficacy in cytotoxic T cells. We created retroviral vectors to deliver ectopic expression of HIF-1ɑ and HIF-2ɑ in mouse CD8+ T cells, together or individually, and with or without sensitivity to their oxygen-dependent inhibitors Von Hippel-Lindau (VHL) and Factor Inhibiting HIF (FIH). We found that HIF-2ɑ, but not HIF-1ɑ, drives broad transcriptional changes in CD8+ T cells, resulting in increased cytotoxic differentiation and cytolytic function against tumor targets. We further found that a specific mutation replacing the hydroxyl group acceptor site for FIH in the HIF-2ɑ isoform gives rise to the most effective anti-tumor T cells after adoptive transfer in vivo. Lastly, we show that co-delivering an FIH-insensitive form of HIF-2ɑ with an anti-CD19 chimeric antigen receptor greatly enhances cytolytic function of human CD8+ T cells against lymphoma cells. These experiments provide a means to increase the anti-tumor efficacy of therapeutic CD8+ T cells via ectopic expression of the HIF transcription factor.

**Graphical abstract:** 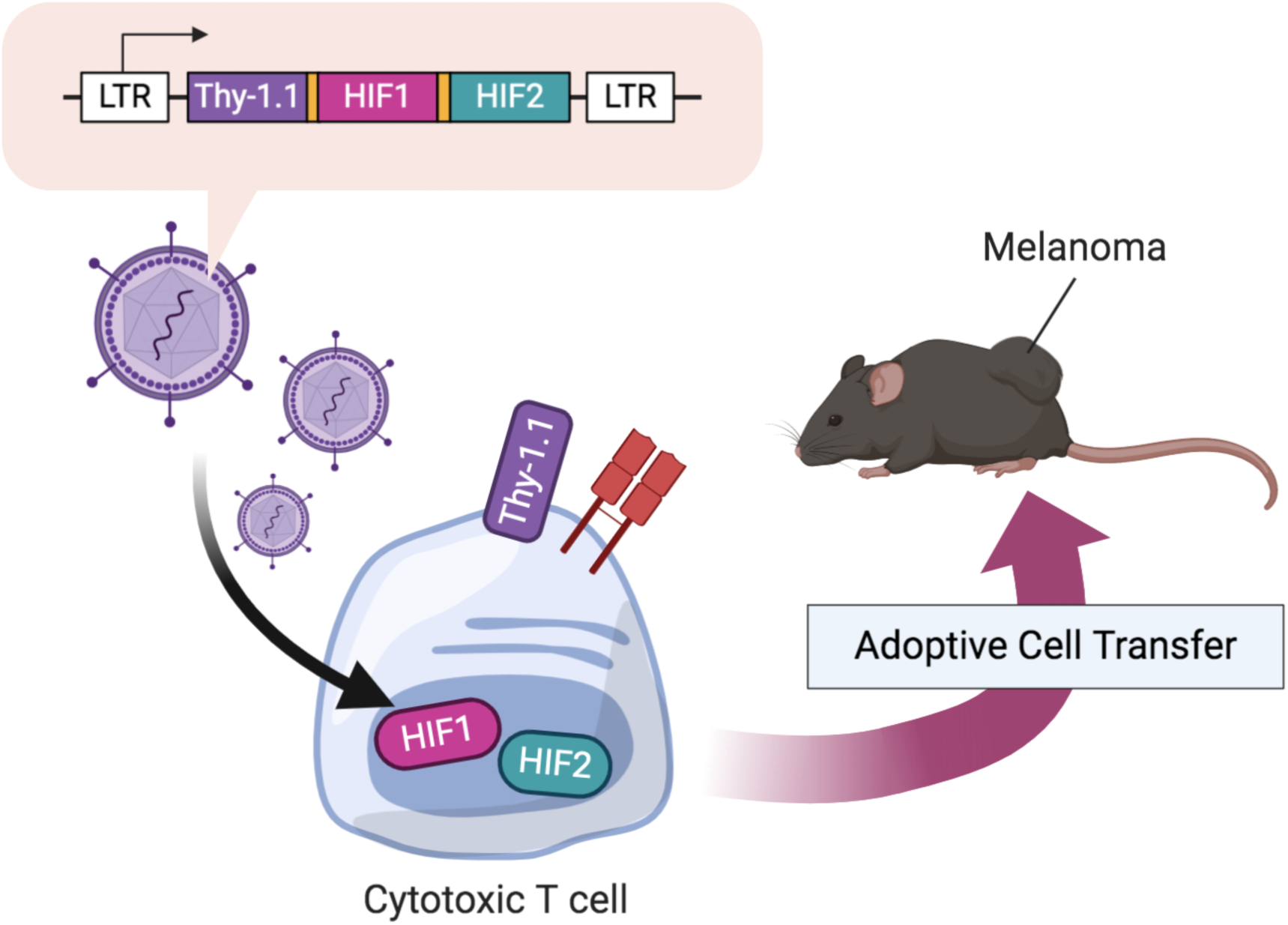

## Introduction

CD8+ T cells are the cytotoxic arm of the adaptive immune system and play an essential role in anti-tumor immunity. One key immunotherapeutic strategy is the redirection of T cells against tumor antigens by retroviral gene transfer of defined T-cell receptors (TCR), or chimeric antigen receptors (CAR), which can be adoptively transferred into cancer patients, and mediate tumor regression in some forms of malignancies (1–3).

The manufacturing process of TCR/CAR T cells can be used to perform additional genetic interventions in order to improve safety and anti-tumor efficacy of the T-cell product. Such interventions include CRISPR-mediated disruption of the endogenous TCR loci (4), disruption of the inhibitor checkpoint protein PD-1 (5) or co-delivery of genes for cytokines (6), chemokines or their receptors (7,8), as well as suicide genes (9). Transcription factors are able to exert large phenotypic changes and guide T-cell differentiation (10) making them an appealing target in TCR/CAR T-cell immunotherapy.

The hypoxia-inducible transcription factors (HIF) are primary regulators of the transcriptional response to hypoxia (11,12). HIFs function as heterodimers composed of an alpha (HIF-1α or HIF-2α) and a beta subunit (ARNT/HIF-1β) and their function is efficiently inhibited by oxygen. In the presence of oxygen, HIF-α subunits are hydroxylated at two conserved prolines by prolyl-hydroxylases (PHD), leading to recognition and ubiquitination by the Von Hippel-Lindau (VHL) protein and resulting in proteasomal degradation (13,14). The second oxygen-sensitive mechanism utilizes the Factor Inhibiting HIF (FIH) enzyme, which hydroxylates a conserved asparagine residue in HIF-α subunits, blocking association with the p300/CBP coactivator and thus HIF transcriptional activity (15,16). These repressive mechanisms are reduced or eliminated in low-oxygen environments leading to HIF stabilization, translocation to the nucleus and induction of a hypoxia-response gene expression program.

Low oxygen availability in tumors and in inflamed tissues has historically been characterized as immunosuppressive (17–19) but a number of recent studies have shown that CD8+ T cells differentiate more efficiently into cytotoxic T cells when cultured in low oxygen (20,21), suggesting that T-cell differentiation is functionally linked to decreased oxygen availability.

This hypothesis was further supported by genetic deletion of VHL (21) or the three PHD isoforms (22) which caused constitutive accumulation of HIF-1α and HIF-2α in T cells and increased cytotoxic differentiation as well as improved rejection of primary and metastatic tumors. Ablation of HIF activity via deletion of ARNT (23) or HIF-1α (24) had the opposite effect, resulting in poorer cytotoxic differentiation, and reduced tumor rejection after adoptive transfer.

Based on these studies, we hypothesized that engineering CD8+ T cells to enhance HIF expression would result in T cells with enhanced anti-tumor properties. To test this hypothesis, we designed retroviral vectors to ectopically express HIF-1α and HIF-2α, alone or in combination, and with or without susceptibility to oxygen-dependent regulation by either VHL- or FIH-dependent mechanisms. We screened these HIF variants in CD8+ T cells to determine whether they enhance *in vivo* anti-tumor efficacy. Our results demonstrate the potential for use of engineered HIF activity as a means to boost T cell function in adoptive T cell immunotherapy.

## Materials and Methods

### Animals

C57BL/6J (CD45.2) animals were purchased from Janvier Labs. Donor TCR-transgenic OT-I mice (catalogue 003831, The Jackson Laboratory) were crossed with mice bearing the CD45.1 congenic marker (catalogue 002014, The Jackson Laboratory) or TdTomato dLck Cre reporter mice (catalogue 007914 and 012837, The Jackson Laboratory).

### Cell lines

HEK293 was a gift from Prof. Dantuma (Karolinska Institute, Stockholm). EL4 was a gift from Prof. H. Stauss (UCL, London). B16-F10 and LLC were purchased from ATCC (CRL-6475 and CRL-1642, respectively). B-luciferase-GFP Raji (B-HCL-010) and B-luciferase K562 (B-HCL-013).

### Vectors

DNA encoding a codon-optimized polycistronic peptide composed of mouse Thy-1.1 (AAR17087.1), mouse HIF-1ɑ (NP_034561.2; P402A, P577A, N813A) and mouse HIF-2ɑ (NP_034267.3; P404A, P530A, N851A) interspersed with picornavirus P2A (GSGATNFSLLKQAGDVEENPGP) and furin (RAKR) cleavage sequences was synthesized by Gene Art (Thermo Fisher). Cell surface and nuclear localization peptides were incorporated into the Thy-1.1 and HIF sequences, respectively. Quick Change II Site-directed mutagenesis (Agilent) was used to revert mutated sites to the native sequence. DNA encoding a codon-optimized polycistronic peptide composed of eGFP (ADQ43426.1), anti-human CD19 CAR and human HIF-2ɑ (NP_001421.2, N847A) interspersed with picornavirus P2A and furin cleavage sequences was synthesized by Gene Art (Thermo Fisher). The coding sequences were cloned into the gamma retroviral vector pMP71, a gift from Christopher Baum (MHH, Hannover). DNA encoding a codon-optimized polycistronic peptide composed of chicken ovalbumin (OVA; P01012.2), eGFP (ADQ43426.1) and neomycin phosphotransferase (NeoR; CAD21956.1) interspersed with P2A and furin cleavage sites was synthesized by Gene Art (Thermo Fisher) and cloned under control of the SV40 promoter in the transposon vector pT2/BH, a gift from Perry Hackett (Addgene plasmid #26556). pCMV-SB11 encoding the sleeping beauty transposase was a gift from Perry Hackett (Addgene plasmid #26552).

### CD8+ T-cell sourcing, activation and restimulation

CD8+ T cells from female and male mice were purified from spleens by CD8ɑ positive selection magnetic bead sorting (Miltenyi) and activated in complete RPMI (Thermo Fisher) supplemented with 55 μM 2-ME (Thermo Fisher) with 2 μg/ml ConA (Sigma) and 10 ng/ml recombinant human IL-7 (R& Systems) for 24 hours before transduction. Whole OT-I splenocytes were activated with 100 ng/ml SIINFEKL (ProImmune) 24 hours before transduction. Transduced CD8+ T cells were restimulated with anti-mouse CD3/CD28 dynabeads (Thermo Fisher) at a 1:1 cell-to-bead ratio or 100 ng/ml SIINFEKL. Transduced CD8+ T cells were expanded in the presence of 10 U/ml recombinant human IL-2 (Sigma) Human CD8+ T cells were purified from donor PBMCs (Cambridge Bioscience or NHSBT) by negative selection magnetic bead sorting (Miltenyi) and activated in complete RPMI supplemented with 30 U/ml IL-2 with anti-human CD3/CD28 dynabeads (Thermo Fisher) at a 1:1 cell-to-bead ratio.

### Retroviral transductions

Sub-confluent HEK293 cultures were transfected with HIF-encoding vectors and helper vector pCL-Eco, a gift from Inder Verma (Addgene plasmid #12371) or pCL-Ampho (Novus). Supernatant media containing retroviral particles was harvested 48 hours after transfection and used fresh or stored at −80°C. Retroviral supernatants were spun onto Retronectin-coated wells (Takara) at 2000 x*g* for 2 hours at 32°C and replaced with activated polyclonal, OT-I, or human CD8+ T cells in fresh RPMI supplemented with 10 or 30 U/ml IL-2. Fresh media was added every 2-3 days. For long-term expansion, transduced cells were re-stimulated weekly with anti-CD3/CD28 dynabeads.

### Flow cytometry

Single cell suspensions were stained with Near-IR Dead Cell Stain Kit (Thermo Fisher) followed by surface, cytoplasmic and/or nuclear staining with the following fluorochrome-labeled monoclonal antibodies: CD45.2 (104), CD3 (145-2C11), 4-1BB (1AH2), CD8ɑ (53-6.7), CD8ɑ (SK1), CD19 (HIB19), CD45RO (UCHL1), CCR7 (3D12), CD45.1 (A20), Vα2 TCR (B20.1), Caspase-3 (active form) (C92-605), LAG3 (C9B7W), CD44 (IM7), PD-1 (J43), IL-2 (JES6-5H4), CD62L (MEL-14), PD-L1 (MIH5), NK-1.1 (PK136), CD147/Basigin (RL73), TCF-1 (S33-966), CTLA-4 (UC10-4F10-11), and IFN-γ (XMG1.2) purchased from BD Biosciences; CD28 (37.51), Fc Block (93), CD45.2 (104), CD80 (16-10A1), NKG2A (16A11), CD3 (17A2), H-2Kb+SIINFEKL (25-D1.16), CD8 (53-6.7), CD122 (5H4), CD45.1 (A20), CD127 (A7R.34), TIM-3 (B8.2C12), ICOS (C398.4A), NKG2D (CX5), CD4 (GK1.5), TCR β chain (H57-597), CD44 (IM7), CD184 (CXCR4) (L276F12), CD27 (LG.3A10), CD62L (MEL-14), OX40 (OX-86), Granzyme B (QA16A02), Perforin (S16009A), Granzyme C (SFC1D8), and IFN-γ (XMG1.2) purchased from BioLegend; 4-1BB (17B5), CD127 (A7R.34), Eomes (Dan11mag), T-bet (eBio4B10), Thy-1.1 (CD90.1) (HIS51), CD44 (IM7), Vβ5.1/5.2 TCR (MR9-4), CD25 (PC61.5), CD4 (RM4-5) purchased from eBioscience and Granzyme B (GB12) purchased from Life Technologies. Staining of cytoplasmic and nuclear antigens was performed using the Fixation/Permeabilization kit (BD Biosciences) and the Transcription Factor buffer set (BD Biosciences), respectively. To measure IFN*γ* secretion cells were incubated overnight in complete RPMI supplemented with 100 ng/ml SIINFEKL with and treated with Golgi Stop (BD Biosciences) 4 hours before intracellular staining and flow cytometry analysis. For proliferation assays, cells were loaded with CellTrace Violet (Thermo Fisher) according to manufacturer’s instructions. Samples were analyzed in a FACSCanto II flow cytometer (BD Biosciences).

### RNA sequencing

OT-I cells were expanded for 5 days in the presence of IL-2 after transduction and sorted on Thy-1.1 surface expression on a BD FACSAria Fusion cell sorter (BD Biosciences) directly into RLT Plus lysis buffer (Qiagen). Total RNA was subjected to quality control with Agilent Tapestation according to the manufacturer’s instructions. To construct libraries suitable for Illumina sequencing the Illumina TruSeq Stranded mRNA Sample preparation protocol which includes cDNA synthesis, ligation of adapters and amplification of indexed libraries was used. The yield and quality of the amplified libraries were analysed using Qubit by Thermo Fisher and the Agilent Tapestation. The indexed cDNA libraries were normalised and combined and the pools were sequenced on the Nextseq 550 for a 50-cycle v2.5 sequencing run generating 75 bp single-end reads. Basecalling and demultiplexing was performed using CASAVA software with default settings generating Fastq files for further downstream mapping and analysis.

### Western blotting

Nuclear protein extracts (15-20 μg) from HEK293 cells transfected with HIF-encoding vectors or from Thy-1.1-purified CD8+ T cells transduced with HIF-encoding vectors were probed with polyclonal antibodies against HIF-1α (NB-100-449 or NB-100-105 Novus Biologicals), HIF-2α (AF2997, R& Systems), Lamin B (sc-6217, Santa Cruz), and Histone 3 (4499S, CST) and detected using infra-red labeled secondary antibodies in an Odyssey imaging system (LI-COR).

### Generation of OVA-expressing cell lines

B16-F10, LLC and EL4 were co-transfected with the transposon vector pT2 encoding OVA, eGFP and neomycin phosphotransferase and the vector encoding transposase SB11. Three days later 400 mg/ml G418 (Gibco) was added to culture media to select for transgene-expressing cells. Successful integration was confirmed by analyzing eGFP fluorescence by flow cytometry. Limiting dilution was used to derive monoclonal OVA-expressing lines for each cell line. OVA presentation was confirmed by flow cytometry using a PE-labeled antibody against surface SIINFEKL bound to H-2Kb (clone 25-D1.16, BioLegend).

### *In vitro* cytotoxicity assay

OVA-expressing GFP-positive LLC or EL4 were mixed with their respective parent cell line in a 1:1 ratio. Transduced OT-I cells were enriched by magnetic bead sorting against Thy-1.1 (Miltenyi) and added to the tumor cells to a final T cell to OVA-positive target ratio of 1:1. Cytotoxicity was assessed by flow cytometry 20 hours later. The ratio of GFP-positive events (target) to GFP-negative tumor cell events (reference) in each test co-culture was divided by the ratio from cultures without OT-I cells to calculate specific cytotoxicity. For, B16-F10-OVA 5000 cells were plated per well in flat-bottom 96 well plates and left to adhere for 1 hour before adding 1000 Thy-1.1+ CD8+ T cells to a final E/T ratio of 1:5. After 20 hours co-culture wells were washed 3 times with PBS to remove T cells and the number of remaining target cells was determined by culturing with 10 μg/ml resazurin (Sigma) and measuring fluorescence signal in a plate reader. Cytotoxicity was calculated relative to wells with no T cells added. To determine cytotoxicity of CAR-transduced human T cells, Raji (GFP+CD19+, target) and K562 (reference) were mixed at 1:1 ratio and co-cultured with varying ratios of GFP+CD8+ T cells. After 20h co-culture cytotoxicity was assessed by flow cytometry and the ratio of target to reference cells used to calculate cytotoxicity. For each donor, cytotoxicity was normalised to the VC level to determine specific cytotoxicity.

### *In vivo* tumor challenge

8 to 12-weeks old female C57BL/6j were inoculated subcutaneously with 5×10^5^ B16-F10-OVA and conditioned 4 days later with peritoneal injection of 300 mg/kg cyclophosphamide (Sigma). On day 8, 0.5-1×10^6^ Thy-1.1-purified transduced OT-I CD8+ T cells were peritoneally injected. Animals were assigned randomly to each experimental group. Tumor volume was measured every 2-3 days with electronic calipers until day 60. Peripheral blood was collected from the tail vein at day 15 and analyzed by flow cytometry. Tumor volume was calculated using the formula 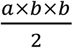 where *a* is the length and *b* is the width of the tumor.

### Tumor-infiltration assessment

8 to 12-weeks old female C57BL/6j were inoculated subcutaneously with 5×10^5^ B16-F10-OVA and conditioned 11 days later with peritoneal injection of 300 mg/kg cyclophosphamide (Sigma). On day 14, 0.5-1×10^6^ Thy-1.1-purified transduced OT-I CD8+ T cells were peritoneally injected. Animals were assigned randomly to each experimental group. On day 19, tumors were dissected, digested with 1 mg/ml Collagenase Type IV (Life Technologies) and 20 U/ml DNAse I (Sigma), and processed in a GentleMACS dissociator (Miltenyi). The tumor single cell suspensions were stained with fluorochrome-labeled antibodies and analysed by flow cytometry.

### Statistics

Statistical analyses were performed with Prism 8 software (GraphPad). Statistical tests used are stated in figure legends. Statistical significance was set at p<0.01 and denoted in figures as *α*.

### Study approval

All animal experiments were approved by the regional animal ethics Committee of Northern Stockholm, Sweden.

## Results

### Generation of retroviral vectors for ectopic expression of HIF-1ɑ and HIF-2ɑ in CD8+ T cells

In order to test whether ectopic expression of HIFs can increase the anti-tumor function of cytotoxic CD8+ T cells we generated retroviral vectors encoding murine HIF-1ɑ and HIF-2ɑ. HIF protein stability and transcriptional activity is regulated by oxygen at the post-translational level (Figure 1A). In the presence of oxygen, prolyl hydroxylases (PHD) hydroxylate conserved proline residues, leading to recognition and ubiquitination by the VHL protein, thus targeting HIFs for proteasomal degradation. FIH hydroxylates a conserved asparagine, which blocks recruitment of the coactivator p300/CBP and impairs HIF transcriptional activity. In order to address the role of HIF regulation we employed site-directed mutagenesis (Figure 1B) to convert conserved proline (P402/P577 in mouse HIF-1α; P405/P530 in mouse HIF-2α) and asparagine (N813 in mouse HIF-1α and N851 in mouse HIF-2α) residues into alanine residues. These alanine residues in the resultant vectors cannot be hydroxylated, and thus prevent proline or asparagine hydroxylation and inhibition of HIF accumulation or activity.

**Figure 1.**
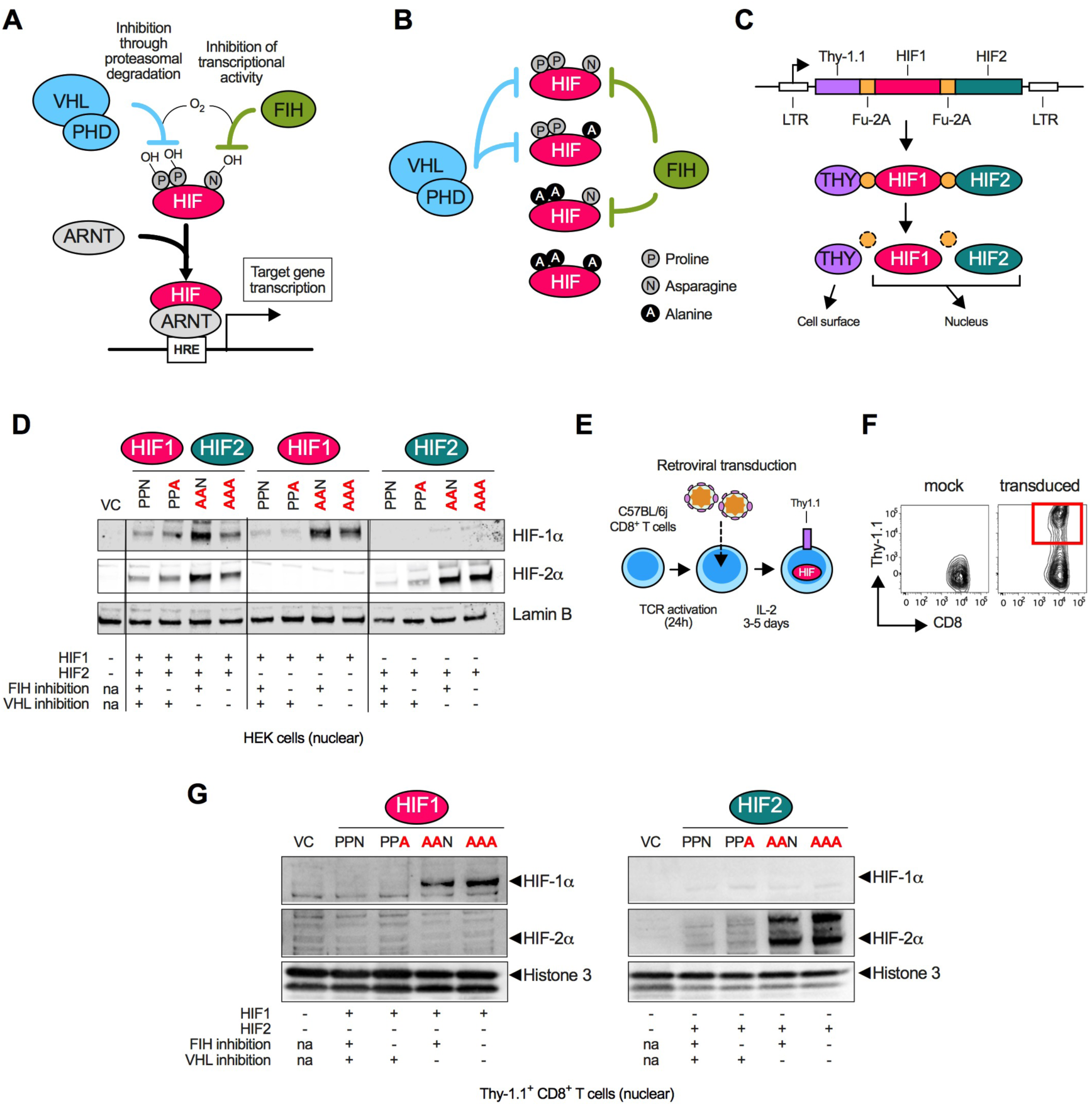
Design and validation of retroviral vectors to drive ectopic expression of HIF proteins in mouse CD8+ T cells. **(A)** HIF transcription factors are post-translationally regulated. Oxygen-dependent hydroxylation at conserved proline (P) residues by proly-hydroxylases (PHD) results in Von Hippel-Lindau (VHL)-mediated proteasomal degradation. Hydroxylation at a conserved asparagine (N) residue by Factor Inhibiting HIF (FIH) prevents recruitment of the coactivator p300/CBP resulting in inhibited transcriptional activity. Once released from VHL/PHD and FIH repression, HIF proteins heterodimerize with Aryl hydrocarbon Receptor Nuclear Translocator (ARNT), translocate to the nucleus, bind hypoxia-responsive elements (HRE) and initiate transcription of target genes. **(B)** Mutation of key amino acid residues modulates HIF regulation. Mutation of conserved prolines (P402, P577 in mouse HIF-1α; P405, P530 in mouse HIF-2α) and of conserved asparagine (N813 in mouse HIF-1α and N851 in mouse HIF-2α) into alanine (A) prevents hydroxylation by PHD and FIH, respectively. **(C)** Retroviral vector design for ectopic HIF expression. After genomic integration the retroviral long terminal repeat (LTR) promoter drives expression of a polycistronic peptide containing Thy-1.1 (THY), HIF-1α (HIF-1ɑ) and HIF-2α (HIF-2ɑ) interspersed with furin cleavage sites and self-cleaving picornavirus 2A sites. Post-translational processing results in separation of the elements. Surface and nuclear localization sequences target Thy-1.1 to the cell surface and HIF isoforms to the nucleus, respectively. **(D)** Nuclear extracts from HEK cells transfected with vectors encoding HIF-1α alone, HIF-2α alone or both probed for mouse HIF-1α, HIF-2α and Lamin B. Vector control (VC) encodes Thy-1.1 alone. **(E)** CD8+ T-cell transduction scheme. Primary CD8+ T cells were purified from mouse (C57BL/6j) splenocytes and activated by T-cell receptor (TCR) triggering for 24 hours before transduction with retroviral particles. Transduced T cells were expanded in the presence of IL-2 for further 3-5 days before subsequent analysis. **(F)** Example of CD8+ T-cell transduction. Representative flow cytometry plot showing retrovirally (RV)-transduced cells expressing Thy-1.1 on the cell surface (red box). **(G)** Nuclear extracts from Thy-1.1+ CD8+ T cells transfected with vectors encoding HIF-1α or HIF-2α probed for mouse HIF-1α, HIF-2α and Histone 3. Vector control (VC) encodes Thy-1.1 alone.

The retroviral vectors designed to carry out these experiments encode a polycistronic peptide composed of Thy-1.1 (transduction marker), HIF-1α, and/or HIF-2α (Figure 1C). These elements are interspersed with furin and self-cleaving picornavirus 2A sites that enable post-translational separation of Thy-1.1 and HIF proteins. Subcellular localization signal peptides traffic Thy-1.1 to the cell surface, and HIFs to the nucleus. The polycistronic nature of the constructs ensures equimolar production of all proteins, while retroviral vector integration in the T-cell genome ensures constitutive and heritable transgene expression.

A library of vectors was generated encoding either both HIF-1ɑ and HIF-2ɑ together, or HIF-1ɑ alone, or HIF-2ɑ alone, and in every mutational arrangement; these are denoted henceforth as PPN (hydroxylated by VHL and FIH), PPA (hydroxylated by VHL only), AAN (hydroxylated by FIH only), and AAA (no hydroxylation by either VHL or FIH). An empty vector control (VC) was generated that encodes Thy-1.1 alone. The ability of these vectors to deliver HIF-1ɑ and HIF-2ɑ protein expression was confirmed in nuclear extracts of transfected HEK cells (Figure 1D). Ectopic expression of HIF isoforms was achieved with every mutational combination. As expected, mutation of proline residues resulted in increased HIF protein accumulation due to the absence of VHL-driven proteasomal degradation.

During retroviral transduction primary mouse CD8+ T cells are activated 24 hours before transduction with retroviral particles, and expanded for a further 3 to 5 days in the presence of IL-2 (Figure 1E). Transduced cells expressing Thy-1.1 on the cell surface can be identified and characterized by flow cytometry (Figure 1F) or purified using magnetic bead sorting for subsequent analysis. Ectopic expression of HIF-1ɑ and HIF-2ɑ was confirmed in nuclear extracts of transduced CD8+ T cells (Figure 1G). Having validated these vectors we next proceeded to investigate the effect of ectopic HIF expression on CD8+ T-cell gene expression, effector differentiation, and anti-tumor function.

### Ectopic HIF-2ɑ expression dramatically alters the CD8+ T-cell transcriptome

To assess the impact of ectopic HIF expression on global gene expression we performed transcriptomic analyses of CD8+ T cells transduced with either HIF-1ɑ or HIF-2ɑ. To increase homogeneity in these analyses we transduced CD8+ T cells from the transgenic OT-I mouse strain in which all T cells express the same MHC class I-restricted TCR specific against a fragment of chicken ovalbumin (OVA) (Figure 2A). Five days after transduction, Thy-1.1^+^ OT-I cells were sorted by flow cytometry for RNA purification followed by RNA sequencing (RNA-seq). In total, 12155 individual transcripts were detected and mapped (Figure 2B). Average transcript frequency and rank distribution was similar to previously published transcriptomic analyses of CD8+ T cells (25,26) including high levels of expression of genes defining CD8+ T-cell identity such as Gzmb, Cd8a, Cd3g, Prf1 and Ifng.

**Figure 2.**
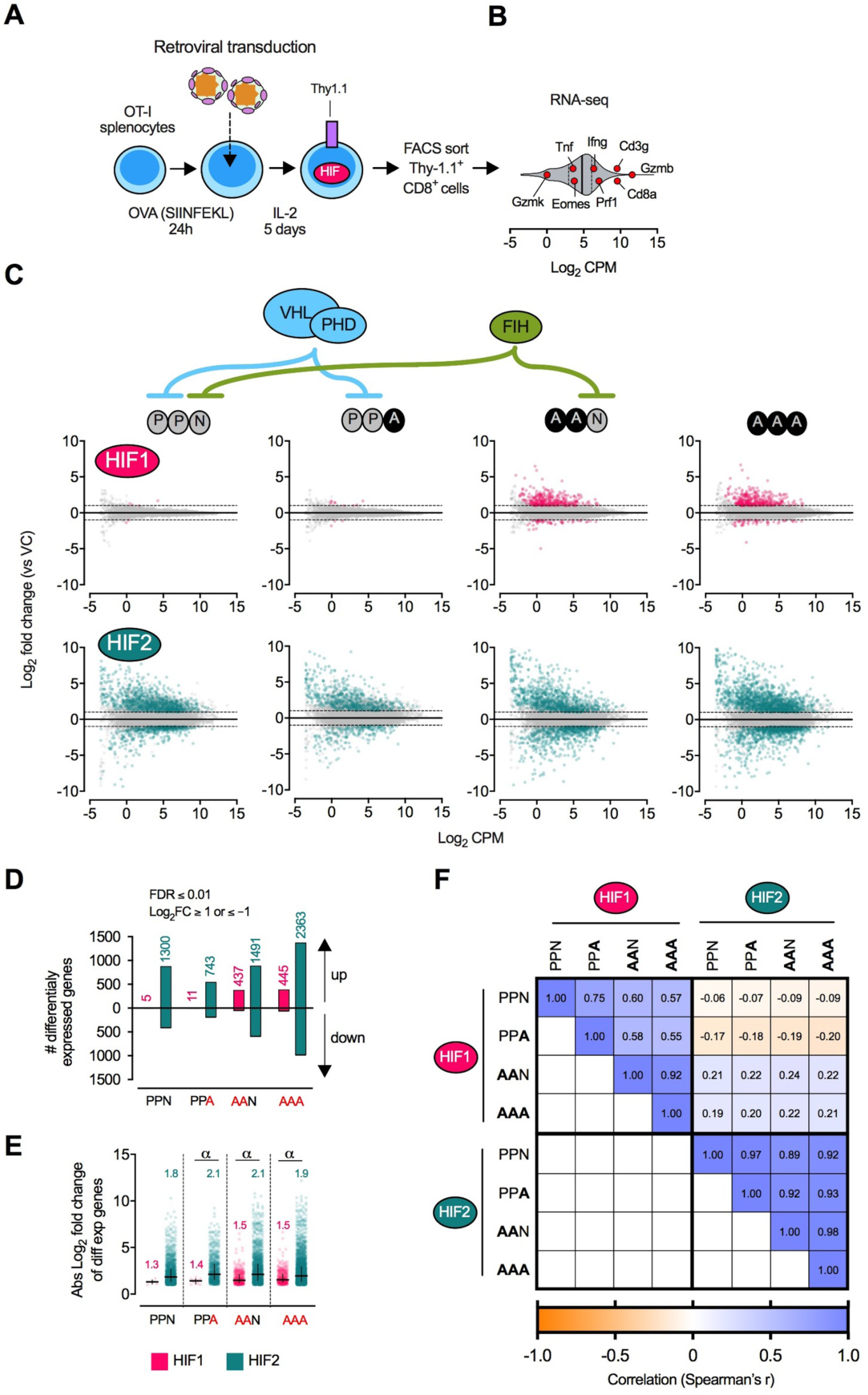
Effects of ectopic expression of HIF-1ɑ or HIF-2ɑ on the CD8+ T-cell transcriptome. **(A)** Ovalbumin (OVA)-specific OT-I splenocytes were activated with an H-2Kb-restricted OVA peptide (SIINFEKL) for 24 hours before transduction with HIF-1ɑ-or HIF-2ɑ-encoding retroviral vectors. After 5 days of expansion in the presence of IL-2, live CD8+ Thy-1.1^+^ cells were sorted by flow cytometry followed by RNA extraction and RNA-seq (n = 3 independent transductions per vector). **(B)** Violin plot representing transcript frequency in Log_2_ counts per million (CPM) of 12155 mapped transcripts. Solid vertical line: median. Dashed vertical line: quartiles. Red circles represent transcripts defining CD8+ T-cell identity. **(C)** Mean-difference plots showing Log_2_ fold change of transcripts in HIF-1ɑ- and HIF-2ɑ-transduced relative to vector control (VC)-transduced CD8+ T cells. Pink and green circles: differentially expressed transcripts as defined by a false discovery rate (FDR) < 0.01 and Log_2_ fold change >1 or <−1. Grey circles: non-differentially expressed transcripts. **(D)** Bar chart summarizing total number of up- and down-regulated transcripts. Values over bars: total number of differentially expressed genes. **(E)** Scattered dot plot showing absolute Log_2_ fold change of differentially expressed transcripts in each transduction. Lines: median and interquartile range. Values over plots: median Log_2_ fold change. α, *P* < 0.01; Kruskal-Wallis with Dunn’s multiple comparison test. **(F)** Heatmap representing correlation in transcript frequency between HIF-1ɑ and HIF-2ɑ-transduced CD8+ T cells. Values in boxes: Spearman’s rank correlation coefficient.

We next compared transcript frequency of HIF-transduced CD8+ T cells with vector control (VC)-transduced CD8+ T cells (Figure 2C). Ectopic expression of VHL-sensitive (PPN and PPA) HIF-1ɑ had minimal impact on gene expression, with only 5 and 11 differentially expressed genes, respectively (Figure 2D). Ablation of VHL control over HIF-1ɑ (AAN and AAA) significantly altered expression of 437 and 445 genes, respectively, suggesting that high levels of HIF-1ɑ protein are required to elicit transcriptional changes in this context. Indeed, we were only able to detect nuclear HIF-1ɑ protein in extracts from CD8+ T cells if VHL regulation was absent (AAN and AAA) (Figure 1G).

Ectopic expression of HIF-2ɑ resulted in a significantly greater number of differentially expressed genes relative to expression of HIF-1ɑ, irrespective of regulatory status (Figure 2C, 2D). HIF-2ɑ altered the expression of 1300, 743, 1491 and 2353 genes in the PPN, PPA, AAN and AAA formats, respectively. Furthermore, the magnitude of changes in gene expression was generally higher in HIF-2ɑ-expressing CD8+ T cells than in those expressing HIF-1ɑ (Figure 2E).

There was high transcriptional similarity amongst cells expressing ectopic HIF-1ɑ, irrespective of susceptibility to VHL or FIH inhibition (Figure 2F). The transcriptional changes elicited by HIF-2ɑ were also similar amongst the PPN, PPA, AAN and AAA variants. However, HIF-1ɑ and HIF-2ɑ appear to alter CD8+ T-cell transcription differently, as shown by low correlation coefficients when comparing differentially expressed genes between the two HIF isoform mutational groups (Figure 2F).

Ectopic HIF-2ɑ altered the expression of several genes involved in CD8+ T-cell function (Figure 3A). Notably, HIF-2ɑ increased expression of perforin (Prf1) and granzyme B (Gzmb), two highly expressed proteins essential for cytolytic function. Ectopic HIF-2ɑ also increased expression of co-stimulators such as CD30 (Tnfrsf8), 4-1BB (Tnfrsf9), OX40 (Tnfrsf4) and ICOS (Icos) and the co-inhibitors LAG3 (Lag3) and CTLA-4 (Ctla4). Reduced expression of interferon-gamma (Ifng), a cytokine that promotes anti-tumor function by inducing antigen presentation in cancer cells, as well as its receptor (Ifngr1 and Ifngr2) was detected in HIF-2ɑ-expressing cells. Expression of the alpha (CD25; Il2ra) and beta chains (CD122; Il2rb) of the IL-2 receptor was increased, suggesting a greater responsiveness to IL-2. Genes involved in T-cell trafficking such as integrins and the bone marrow-homing receptor CXCR4 (Cxcr4) (27) were also induced by ectopic HIF-2ɑ expression.

**Figure 3.**
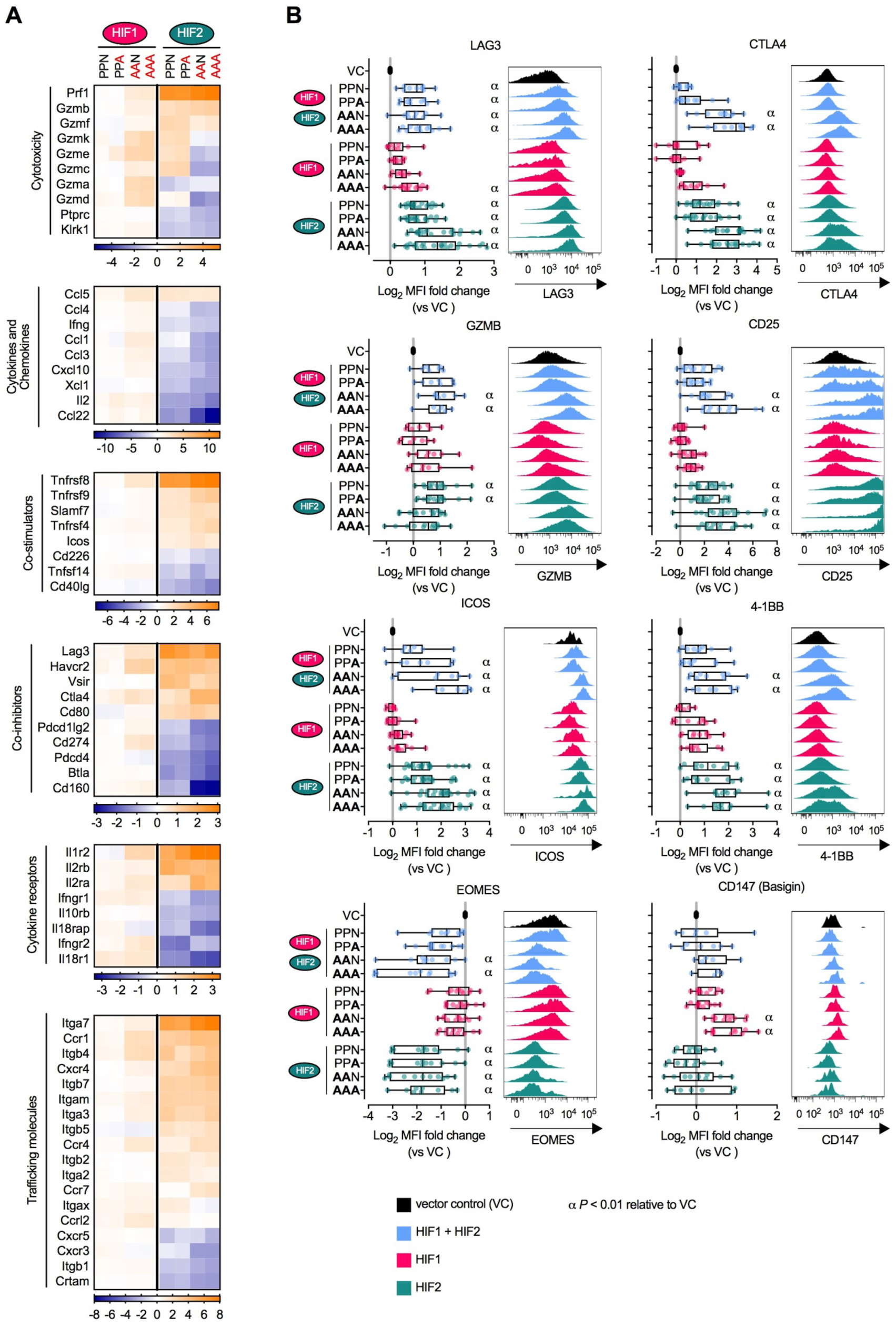
Effects of ectopic expression of HIF-1ɑ or HIF-2ɑ in genes involved in CD8+ T-cell function. **(A)** Heatmaps showing Log_2_ fold change of transcripts involved in functional aspects of CD8+ T cells. **(B)** Expression of differentiation markers determined by flow cytometry in CD8+ T cells transduced with vectors encoding HIF-1ɑ and HIF-2ɑ, HIF-1ɑ alone or HIF-2ɑ alone (day 3 to 5 post-transduction). Data expressed as Log_2_ fold change of median fluorescence intensity (MFI) relative to VC-transduced cells. Each data point represents an independent transduction (n=4-24). Results are pooled from a minimum of two independent experiments. α, *P* < 0.01; one-way ANOVA with Dunnett’s multiple comparison test relative to VC. Histograms are representative flow cytometry plots for each parameter and are pre-gated on live, singlet, CD8+ Thy-1.1+ events.

The broad transcriptional changes induced by HIF-2ɑ can be partially explained by altered expression of a wide array of transcriptional factors and transcriptional modulators (Supplementary Figure 1A and B). While few transcriptional modulators were altered by HIF-1ɑ, ectopic expression of HIF-2ɑ significantly altered expression of 77, 40, 98, and 135 transcriptional modulator genes in the PPN, PPA, AAN and AAA mutational variants respectively, thus amplifying the transcriptional impact of HIF-2ɑ. Notably, HIF-2ɑ decreased expression of transcription factors associated with naive/memory as well as the exhausted state (Tcf7 (28,29), Tox (30), Nr4a1, Nr4a3 (31) and Eomes (32)). Transcripts associated with a response to hypoxia (Supplementary Figure 1C) and induction of glycolysis (Supplementary Figure 1D) were also upregulated in CD8+ T cells expressing HIF-2ɑ, but not in those expressing HIF-1ɑ.

### HIF-2ɑ drives phenotypical changes in CD8+ T cells, particularly in the absence of VHL suppression

We next sought to confirm if the transcriptional changes induced by ectopic HIF expression translated to alterations in protein expression. For that we transduced CD8+ T cells with vectors encoding HIF-1ɑ and HIF-2ɑ together, HIF-1ɑ alone, or HIF-2ɑ alone and analysed their phenotype by flow cytometry (Figure 3B). In line with the RNA-seq results described above, expression of the co-inhibitors LAG3 and CTLA-4 was augmented by HIF-2ɑ, particularly in the absence of VHL inhibition (AAN and AAA). Ectopic co-expression of HIF-1ɑ and HIF-2ɑ yielded similar results to HIF-2ɑ alone. Increased expression of the IL-2 receptor chains CD25 (Figure 3B) and CD122 (Supplementary Figure 2A) was also confirmed at the protein level as was expression of the chemokine receptor CXCR4 (Supplementary Figure 2A). Likewise, expression of GZMB, ICOS and 4-1BB, key components of the cytotoxic function of CD8+ T cells, was increased by ectopic expression of HIF-2ɑ (Figure 3B). Notably, GZMB expression was only significantly increased when HIF-2ɑ was susceptible to VHL regulation (PPN and PPA). Suppression of the transcription factor EOMES by HIF-2ɑ was also confirmed by intranuclear flow cytometry. Of the assayed proteins only CD147 (Basigin), a known direct HIF-1ɑ target (33), was augmented by HIF-1ɑ (Figure 3B). The increased expression of GZMB, LAG3, CTLA-4, CD25 and 4-1BB and suppression of EOMES was confirmed in CD8+ T cells transduced with HIF-2ɑ and cultured over a period of 21 days with weekly restimulation (Supplementary Figure 2B and C), confirming the long-term stability of the phenotypes conferred by the integrated retroviral vectors. Altogether, these data show that ectopic HIF-2ɑ expression, but not HIF-1ɑ, was able to substantially alter the phenotype of CD8+ T cells and that the magnitude of those changes is greater in the absence of inhibition by VHL.

### Ectopic expression of VHL-insensitive HIF-2ɑ reduces cell proliferation, TCR expression and IFNγ secretion while VHL-sensitive HIF-2ɑ enhances CD8+ T-cell cytotoxicity against cancer cells

During long-term culture of HIF-transduced CD8+ T cells (Supplementary Figure 2B) the frequency of cells expressing VHL-insensitive HIF-2ɑ (AAN and AAA) diminished progressively over time (Figure 4A) unlike the remaining transduced populations whose frequency in culture increased slightly over 21 days. This was due to reduced proliferation of VHL-insensitive HIF-2ɑ-transduced CD8+ T cells (Figure 4B, 4C and Supplementary Figure 3A) and not due to increased apoptosis (Supplementary Figure 3B).

**Figure 4.**
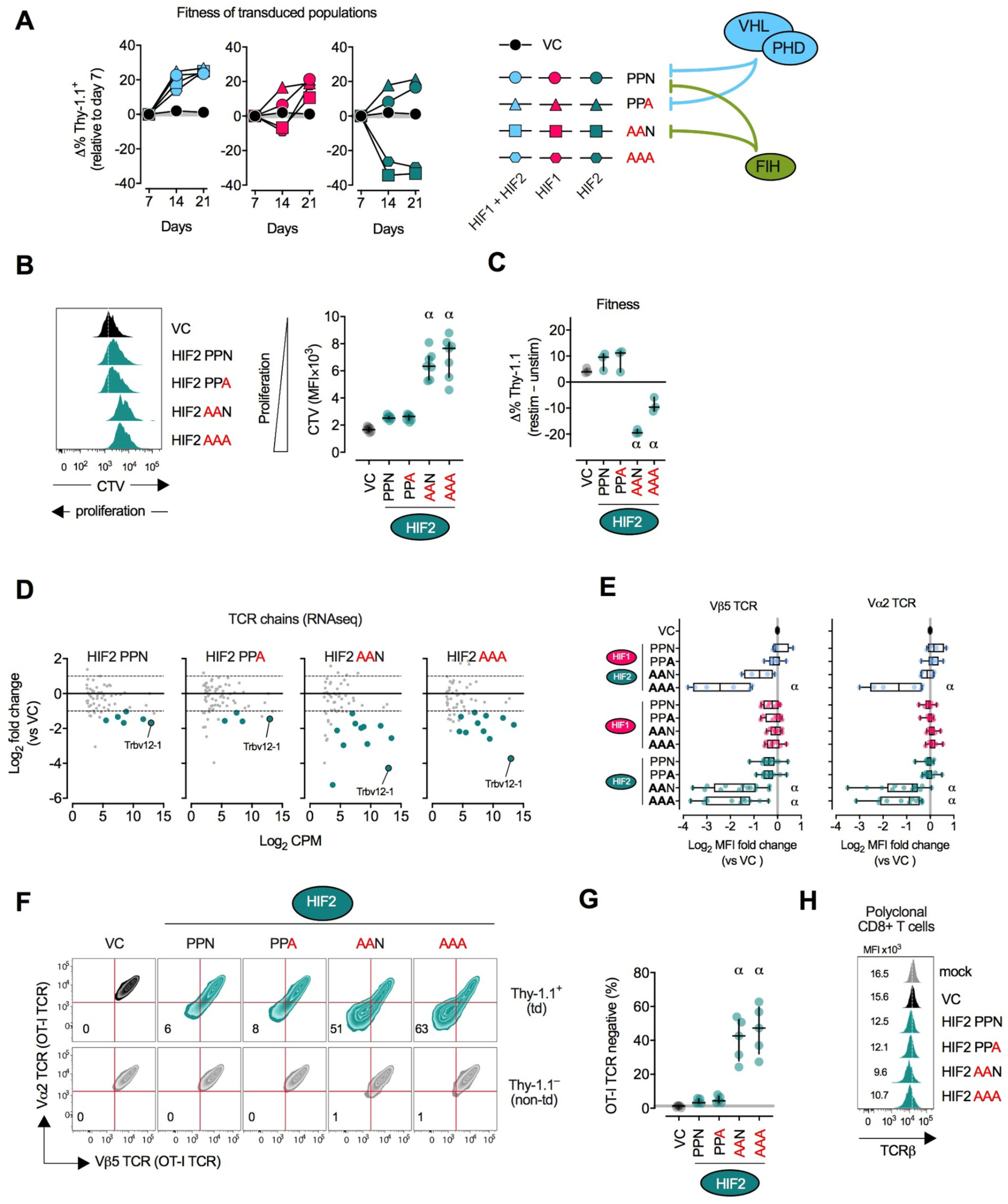
Ectopic expression of VHL-insensitive HIF-2ɑ reduces CD8+ T-cell proliferation and drives TCR loss. **(A)** Fitness of HIF-transduced CD8+ T cells over time. CD8+ T cells were transduced with HIF-1ɑ and HIF-2ɑ-coding vectors and cultured for 21 days in the presence of IL-2. Cells were restimulated with CD3/CD28 beads on days 7 and 14. Fitness was calculated as the difference in % of Thy-1.1^+^ cells in culture relative to day 7 (Δ% Thy-1.1^+^). VC: vector control. **(B)** Proliferation of HIF-2ɑ-transduced CD8+ T cells. Cells were loaded with CellTrace Violet (CTV) proliferation dye 6 days after transduction and were restimulated with αCD3/CD28 beads for 3 days. Proliferation was determined by CTV dilution in flow cytometry. Left: representative histograms pre-gated on live, singlet, CD8+. Thy-1.1+ events. Right: summary data showing CTV mean fluorescence intensity (MFI). n = 7 independent transductions. Lines: median and interquartile range. **(C)** Fitness of HIF-2ɑ-transduced cells after restimulation. Fitness was calculated as the difference in % of Thy-1.1^+^ cells in culture between restimulated and unstimulated cultures (Δ% Thy-1.1^+^). n = 3 independent transductions. Lines: median and interquartile range. **(D)** Mean-difference plots showing Log_2_ fold change of TCR chain-coding transcripts in HIF-2ɑ-transduced relative to VC-transduced CD8+ T cells. Green circles: differentially expressed transcripts as defined by a false discovery rate (FDR) < 0.01 and Log_2_ fold change >1 or <−1. Grey circles: non-differentially expressed transcripts. Trbv12-1 codes the Vβ5 segment of the TCRβ chain. **(E)** Expression of TCR Vα2 and TCR Vβ5 chains determined by flow cytometry in OT-I CD8+ T cells transduced with vectors encoding HIF-1ɑ and HIF-2ɑ, HIF-1ɑ alone or HIF-2ɑ alone (day 3 to 5 post-transduction). Data expressed as Log_2_ fold change of MFI relative to VC-transduced cells. Each data point represents an independent transduction (n=4-24). Results are pooled from a minimum of two independent experiments. **(F)** Surface expression of OT-I TCR chains in HIF-2ɑ-transduced OT-I cells on day 4 post-transduction. Flow cytometry zebra plots pre-gated on live, singlet, CD8+. events showing surface expression of OT-I TCR Vα2 and TCR Vβ5 chains in transduced (Thy-1.1^+^; top row) and non-transduced (Thy-1.1−; bottom row). Values are percentage within the double-negative quadrant. **(G)** Frequency of TCR-negative cells. n=5 independent transductions. Lines: median and interquartile range. Grey horizontal line: median of VC group. **(H)** Surface expression of the TCRβ constant chain in polyclonal CD8+ T cells transduced with HIF-2ɑ-coding vectors. Values in histograms are MFI x10^3^. α, *P* < 0.01; one-way ANOVA with Dunnett’s multiple comparison test relative to VC.

Transcriptomic analysis revealed downregulation of genes encoding TCR chains in HIF-2ɑ-transduced cells, particularly when VHL regulation was absent (Figure 4D). OT-I cells transduced with VHL-insensitive (AAN and AAA) HIF-2ɑ had reduced surface expression of TCR alpha (Vɑ2) and TCR beta (Vβ5) chains (Figure 4E), while VHL-sensitive HIF-2ɑ or any form of HIF-1ɑ had no effect on TCR expression. Loss of TCR chains occurred synchronously resulting in TCR loss in over 50% of AAN and AAA HIF-2ɑ-expressing cells (Figure 4F and 4G) TCR loss occurred progressively after transduction (Supplementary Figure 3C) and was also observed in polyclonal CD8+ T cells expressing these vectors (Figure 4H).

Production of IFNγ after TCR triggering with OVA was also reduced in OT-I cells transduced with VHL-insensitive HIF-2ɑ (AAN and AAA) (Figure 5A and 5B), a likely consequence of reduced TCR expression, since stimulation with PMA and ionomycin, which bypasses the TCR, resulted in unchanged IFNγ secretion (Supplementary Figure 2C).

**Figure 5.**
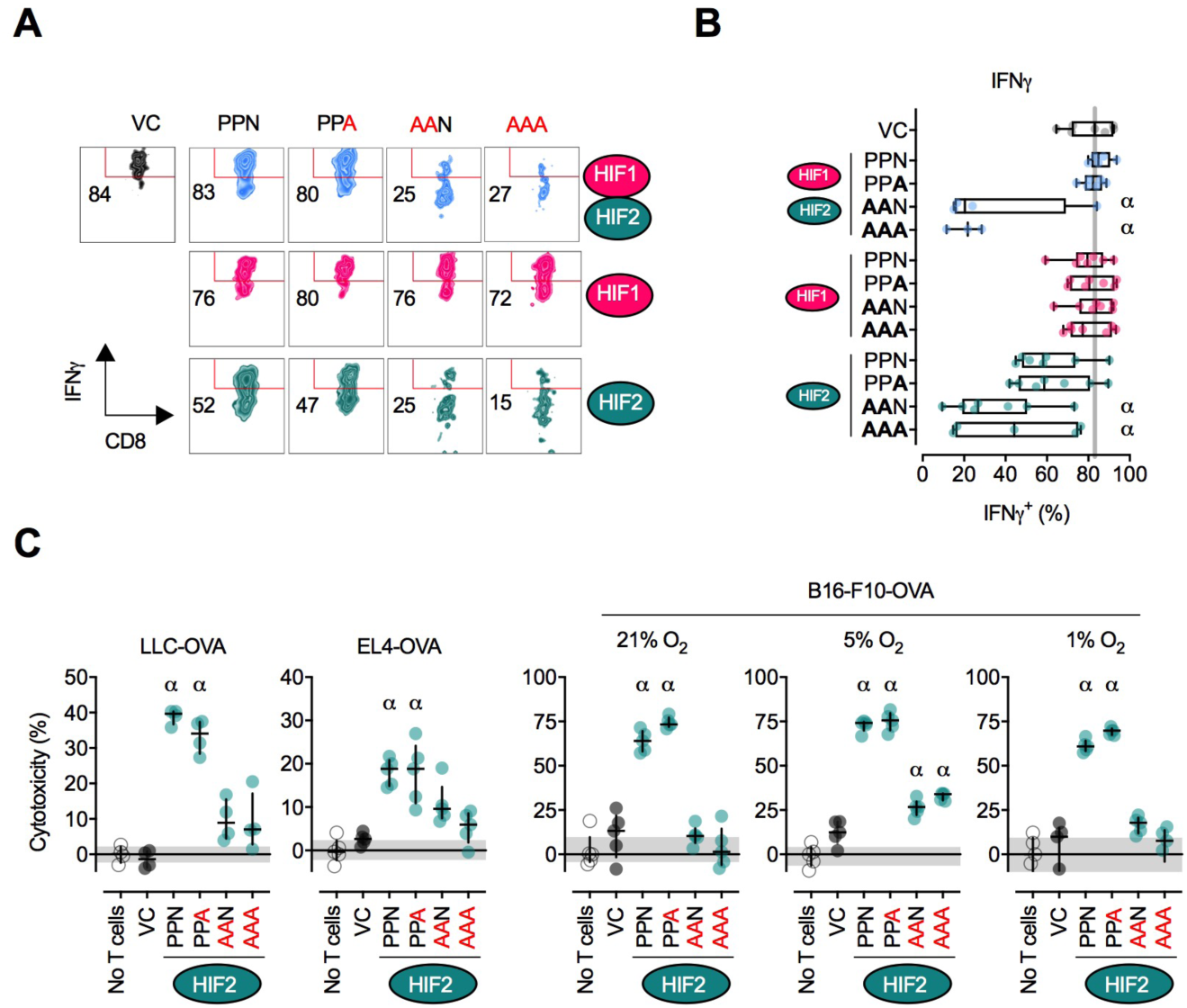
Ectopic expression of HIF-2ɑ enhances cytotoxicity against tumor cells. **(A)** Interferon-γ (IFNγ) secretion determined by intracellular cytokine flow cytometry in OT-I CD8+ T cells transduced with vectors encoding HIF-1ɑ and HIF-2ɑ, HIF-1ɑ alone or HIF-2ɑ alone and restimulated for 4 hours with 1 μM OVA (SIINFEKL) peptide. Values are the percentage within the IFNγ^+^ gate. Pre-gated on live, singlet, CD8+. Thy-1.1+ events **(B)** Summary data expressed as % IFNγ^+^ cells. Each data point represents an independent transduction (n=3-7). Results are pooled from a minimum of two independent experiments. **(C)** OT-I CD8+ T cells transduced with HIF-2ɑ-coding vectors were Thy-1.1-enriched and co-cultured with OVA-expressing EL4 thymoma (EL4-OVA) or Lewis Lung Carcinoma (LLC-OVA) or B16-F10 melanoma (B16-F10-OVA) tumor cells at an effector:target ratio of 1:1 (EL4, LLC) or 1:5 (B16-F10). Cytotoxicity was assessed after 20 hours of co-culture at 21%, 5% or 1% O_2_. *n*=4-5 technical replicates. Lines: median and interquartile range. Grey horizontal area: interquartile range of no T cell control. α, *P* < 0.01; one-way ANOVA with Dunnett’s multiple comparison test relative to VC.

To assess how the phenotypic and functional changes caused by ectopic HIF-2ɑ expression affect cytotoxicity against tumor cells, we co-cultured transduced OT-I cells with OVA-expressing EL4 thymoma, Lewis Lung Carcinoma (LLC) or B16-F10 melanoma cancer cells (Figure 5C). Ectopic HIF-2ɑ expression increased cytotoxicity against EL4, LLC and B16-F10 cells, but only if sensitivity to VHL was retained (PPN and PPA). Expression of VHL-insensitive HIF-2ɑ (AAN and AAA) had no effect on cytotoxicity. Killing of B16-F10 cells was similar whether co-culture occurred at atmospheric oxygen conditions (21% O_2_) or at physiologically relevant oxygen tensions (5% and 1% O_2_). This result can be partly attributed to increased GZMB levels (Figure 3B) and to largely unchanged proliferation rates, TCR expression and IFNγ secretion in the HIF-2ɑ PPN and PPA groups (Figures 4 and 5). Overall, these results reveal that ectopic expression of highly stable HIF-2ɑ protein (AAN and AAA) in CD8+ T cells leads to a deleterious phenotype while expression VHL-sensitive (PPN and PPA) HIF-2ɑ potentiates *in vitro* cytotoxicity against tumor cells.

### Adoptive transfer of CD8+ T cells ectopically expressing FIH-insensitive HIF-2ɑ results in delayed tumor growth and increased survival rate

In therapy, adoptively transferred CAR/TCR engineered T cells must engraft, proliferate, traffic to tumor sites and eliminate cancer cells in order to deliver therapeutic benefit. To test the anti-tumor function of HIF-transduced CD8+ T cells we employed a mouse model of adoptive cell therapy (ACT) (Figure 6A). Wild-type mice were inoculated with OVA-expressing B16-F10 melanoma cells, followed by lymphodepleting chemotherapy and adoptive transfer of HIF-1ɑ- or HIF-2ɑ-transduced OT-I cells (Supplementary Figure 4A). A week after T-cell injection, peripheral blood was harvested, and the frequency of transduced OT-I cells assessed by flow cytometry (Figure 6B and Supplementary Figure 4B). The engraftment of T cells expressing VHL- and FIH-sensitive (PPN) HIF-2ɑ was significantly increased, while cells expressing VHL-insensitive (AAN and AAA) HIF-2ɑ were nearly absent in peripheral blood. The frequency of T cells expressing VHL- and FIH-insensitive (AAA) HIF-1ɑ was also reduced relative to the control group.

**Figure 6.**
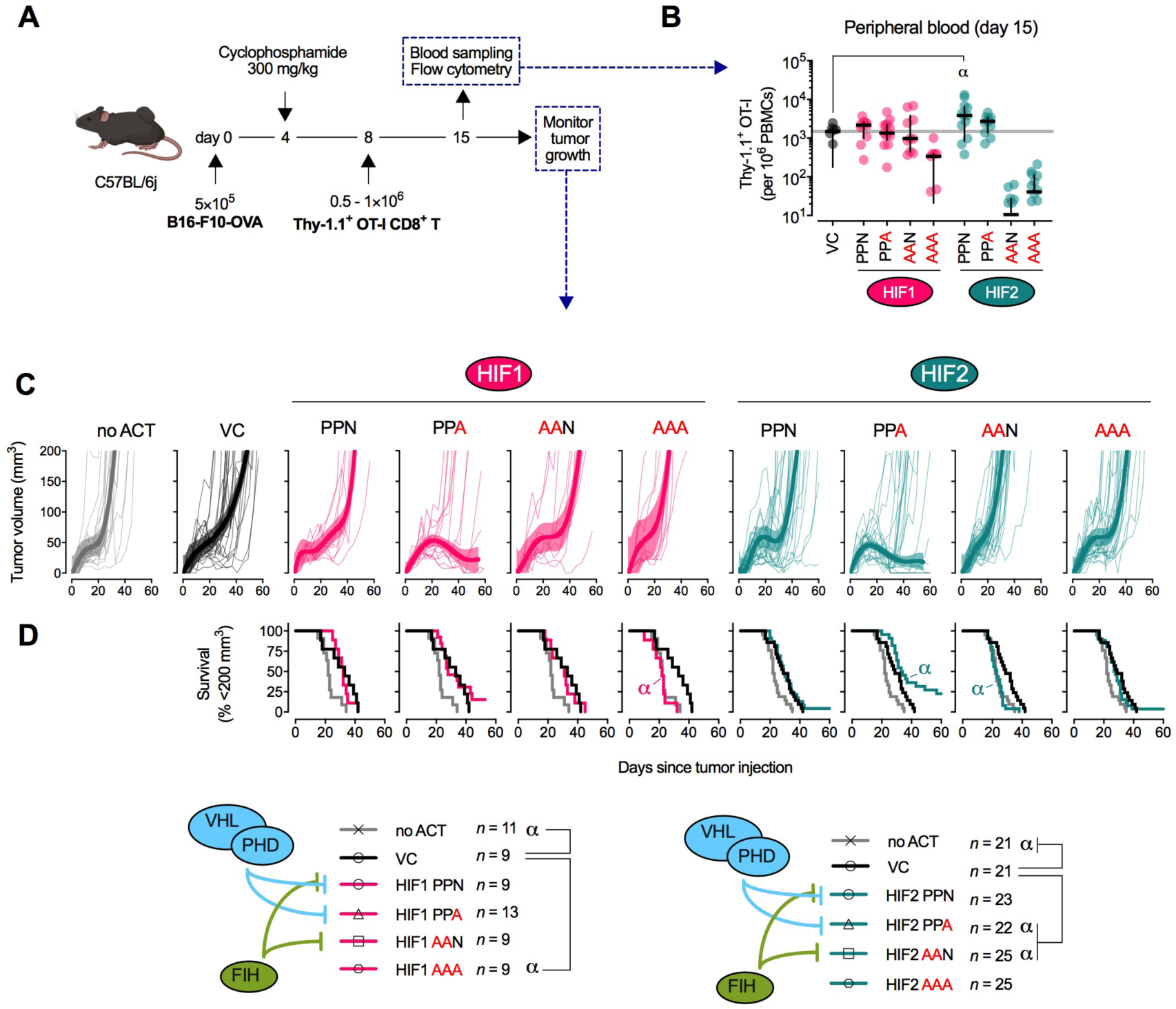
Adoptive transfer of CD8+ T cells ectopically expressing FIH-insensitive HIF-2ɑ results in delayed tumor growth and increased survival rate. **(A)** Adoptive cell therapy (ACT) model. C57BL/6j mice were injected subcutaneously with 5×10^5^ OVA-expressing B16-F10 (B16-F10-OVA) and 4 days later were lymphodepleted with 300 mg/kg cyclophosphamide. On day 8, 0.5-1×10^6^ HIF-transduced (Thy-1.1 enriched) OVA-specific OT-I CD8+ T cells were adoptively transferred into tumor-bearing mice. Peripheral blood was sampled at day 15 and analysed by flow cytometry. Tumor growth was monitored every 2-3 days until day 60. **(B)** Frequency of HIF-transduced OT-I cells per million PBMCs in peripheral blood. n = 8-13 animals pooled from two independent experiments. Grey horizontal line: median of VC group. α, *P* < 0.01; one-way ANOVA with Dunnett’s multiple comparison test relative to VC. **(C)** B16-F10-OVA tumor growth after ACT. B16-F10-OVA tumor volume measured until day 60 after mice received VC-HIF-1ɑ- or HIF-2ɑ-transduced OT-I cells on day 8. Thin lines: tumor growth from individual animals. Thick line: centred sixth order polynomial curve. Shaded area: 99% confidence level interval. *n* = 9-25 animals per group pooled from two (HIF-1ɑ) or four (HIF-2ɑ) independent experiments. **(D)** Survival curves for tumor growth shown in (D). Threshold for survival was set at 200 mm^3^. Grey line: no ACT. Black line: ACT of VC-transduced OT-I. Pink or green lines: ACT of HIF-1ɑ- or HIF-2ɑ-transduced OT-I, respectively. α, *P* < 0.01; log-rank (Mantel-Cox) test relative to VC.

In this model, transfer of VC-transduced OT-I CD8+ T cells delayed tumor growth relative to the untreated (no ACT) group, and increased median survival from 22 to 29 days. However, there were no complete regressions in this group (Figure 6D and Supplementary Figure 4C). Adoptive transfer of T cells expressing VHL-sensitive, but FIH-insensitive (PPA) HIF-2ɑ both significantly delayed tumor growth while boosting median survival to 34 days, and fully eliminated tumors in 23% of the treated animals (Figure 6D and Supplementary Figure 4C). In contrast, animals receiving T cells expressing VHL-insensitive but FIH-sensitive (AAN) HIF-2ɑ performed significantly worse than controls, resulting in median survival times comparable to those of the untreated group (23 days). Likewise, T cells expressing VHL- and FIH-insensitive (AAA) HIF-1ɑ failed to protect animals from tumor growth, while the FIH-insensitive (PPA) HIF-1ɑ variant resulted in complete tumor clearance in 15% of animals; albeit without significantly delaying tumor growth (Figure 6D and Supplementary Figure 4C). These tumor rejection rates did not correlate with a differential ability to infiltrate tumor sites, since cells expressing VHL-sensitive HIF-2ɑ (PPN and PPA) infiltrated tumors at the same rate as controls, while cells expressing VHL-insensitive HIF-2ɑ (AAN and AAA) failed to infiltrate the tumor site (Supplementary Figure 4D-F). Infiltration of endogenous CD8+ T cells was unaltered (Supplementary Figure 4G). These experiments reveal an unexpected role of the FIH hydroxylation site on the HIF-2ɑ protein for the *in vivo* anti-tumor function of CD8+ T cells, and indicate an important role for oxygen-mediated transcriptional control of HIF-2a that is qualitatively different from the role played by the VHL pathway.

### Human CD8+ T cells co-expressing a CAR and FIH-insensitive HIF-2ɑ have increased cytolytic activity against lymphoma cells

To test whether HIF-2ɑ expression could improve the anti-tumor function of human therapeutic T cells we designed a vector encoding an anti-CD19 CAR alone or in combination with FIH-insensitive HIF-2ɑ (Figure 7A and Supplementary Figure 5A). CD8+ T cells purified from multiple human PBMC donors were successfully transduced with CAR vectors as measured by GFP expression (Figure 7B) and consistently showed increased CCR7 levels (Supplementary Figure 5B) in agreement with the RNAseq analysis of mouse CD8+ T cells (Figure 3A). Cytolytic function of CAR-transduced CD8+ T cells against Raji (CD19+) lymphoma cells was significantly increased when HIF-2ɑ was co-expressed (Figure 7C and Supplementary Figure 5C). This result replicates the observations in mouse cells and further supports the translational potential of ectopic expression of HIF-2ɑ in improving the anti-tumor efficacy of T cell products.

**Figure 7.**
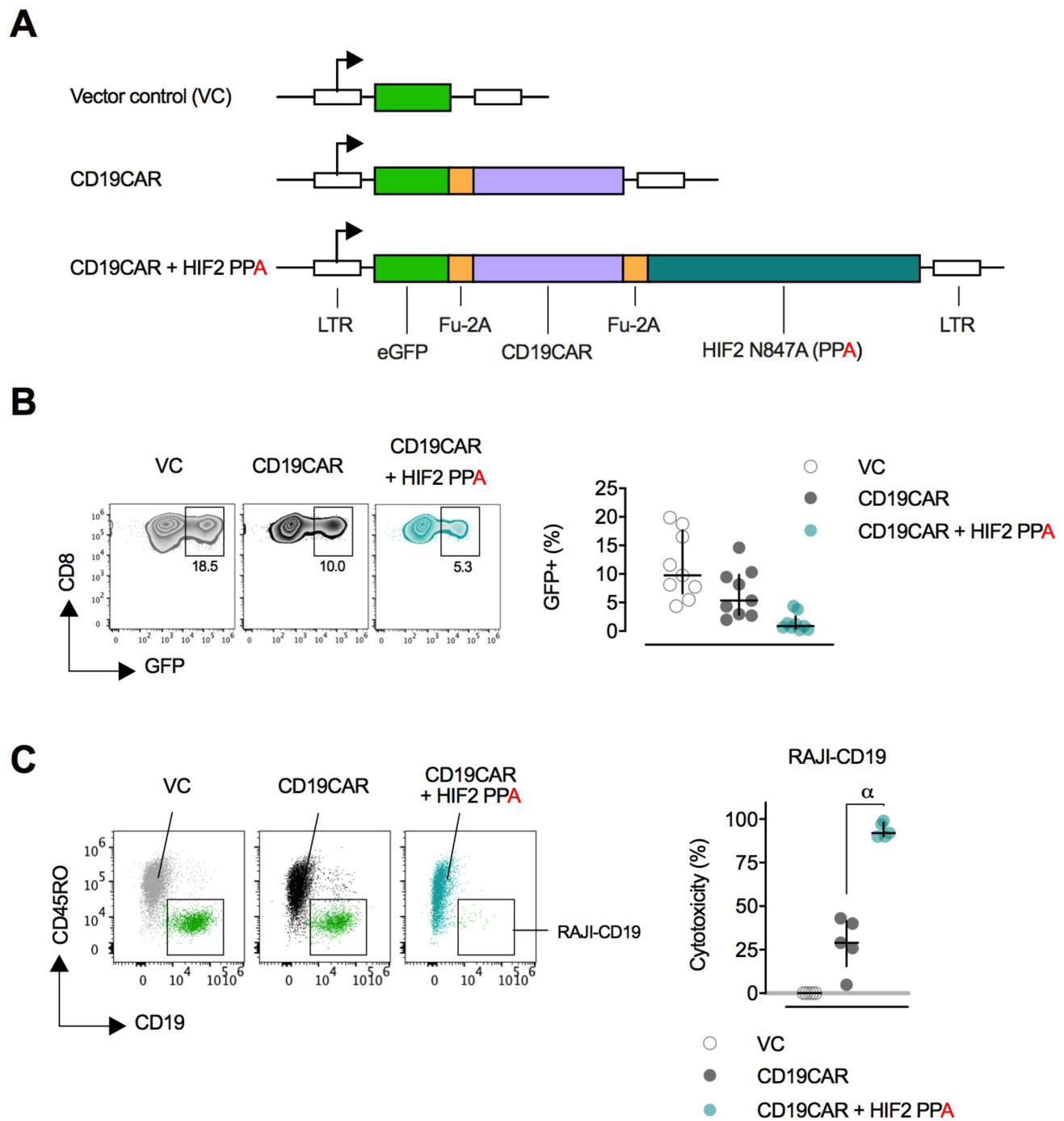
FIH-insensitive HIF-2ɑ increases cytotoxicity of human CD8+ CAR-T cells against lymphoma cells. **(A)** Retroviral vector design for co-expression of an anti-CD19 CAR and FIH-insensitive (N847A; PPA) human HIF-2ɑ. LTR: long terminal repeats. Fu-2A: furin and picornavirus 2A self-cleaving sequence. **(B)** Representative flow cytometry plots of transduced (GFP+) human CD8+ T cells pre-gated on live, singlet, CD8+ events and summary data of transduction efficiency of 9 donors. Lines: median and interquartile range. **(C)** Cytotoxicity of VC- or CAR-transduced human CD8+ T cells against GFP+ CD19+ RAJI lymphoma cells after 20 hours of co-culture at a 1:1 effector-to-target ratio. Representative flow cytometry showing transduced (GFP+, CD45RO+) CD8+ T cells and RAJI targets (GFP+, CD19+) after gating on live, singlet GFP+ events. Summary data of specific cytotoxicity of 5 donors. Lines: median and interquartile range. α, *P* < 0.01; one-way ANOVA with Dunnett’s multiple comparison test relative to VC.

## Discussion

In this study we set out to determine if ectopic HIF expression can be used to increase the anti-tumor efficacy of therapeutic CD8+ T cells with the ultimate goal of improving adoptive T-cell therapy protocols for cancer. Retroviral vectors are currently used to deliver ectopic expression of cancer antigen-specific TCRs or CARs to T cells and can be easily modified to deliver additional proteins (7,8). Previous studies employing T-cell specific deletion of VHL (21), PHDs (22), ARNT (23), HIF-1ɑ, and HIF-2ɑ (24) or hypoxic-priming (20) of CD8+ T cells laid the groundwork for this study by showing that HIF activity drives cytotoxic differentiation and *in vivo* anti-tumor function.

Here we took an agnostic approach regarding which of the two main HIF isoforms and which oxygen-dependent suppression mechanisms could most potently boost anti-tumor effects by T cells. Previous studies would place non-degradable HIF-1ɑ as the strongest candidate for over-expression, given that VHL- and PHD-null T cells reject tumors more efficiently(21,22), while HIF-1ɑ-null T cells perform quite poorly in immunotherapy models(24).

Our findings reveal that in an ectopic expression setting it is HIF-2ɑ, and not HIF-1ɑ, that is able to most potently boost CD8+ T-cell anti-tumor cytotoxicity. Ectopic HIF-2ɑ was able to elicit broad changes in gene expression by altering the T cell transcription factor network, irrespective of VHL and FIH suppression. However, the magnitude of change in gene expression was typically higher when HIF-2ɑ was resistant to VHL-mediated proteolytic degradation. The transcripts altered by HIF-2ɑ extend beyond classical HIF target genes. This is likely due to altered expression of a network of transcriptional modulators, many of them known to have implications for CD8+ T-cell differentiation, including *Tcf7, Nr4a1, Nr4a3, Tox* and *Eomes*, all of which drive memory and/or exhaustion phenotypes, and are downregulated by HIF-2ɑ. Ectopic HIF-2ɑ expression does not result in a gene signature that clearly identifies a memory, effector or exhausted population, generating instead an unconventional phenotype with upregulation of co-stimulators (4-1BB, ICOS), effector proteins (GZMB, Perforin), but also co-inhibitors (CTLA-4, LAG3) and downregulation of IFN*γ*.

While transcriptional changes detected by RNAseq were largely replicated at protein level, it was at the functional level that oxygen-dependent regulation of HIF-2ɑ proved to be most important. Proteolytic-resistant HIF-2ɑ (AAN and AAA) severely impaired T cell function by restricting proliferation, causing TCR downregulation and decreased IFNγ secretion. These results are surprising given the greater anti-tumor function of VHL- and PHD-null T cells reported previously (21,22). Co-culture with a variety of tumor targets consistently revealed that CD8+ T cells expressing VHL-sensitive (PPN and PPA) HIF-2ɑ had the highest cytolytic function suggesting that intermediate levels of HIF-2ɑ protein offer the most benefit.

Regulation of HIF-2ɑ has striking effects in the adoptive T cell therapy models we employed. When transferred to melanoma-bearing mice, CD8+ T cells expressing HIF-2ɑ proteins that are insensitive to FIH, but sensitive to VHL, provide the best protection against tumor growth. The exact opposite arrangement, FIH-sensitive and VHL-insensitive HIF-2ɑ, results in a complete loss of tumor protection by T cells. This role for FIH hydroxylation was only revealed in an *in vivo* setting. Transcriptional analysis showed great similarity between the PPN and PPA groups (0.97 Spearman r) and between the AAN and AAA groups (0.98 Spearman r) despite hydroxylation at N851 being reported as crucial for the transcriptional activity of HIF-2ɑ. It is possible that the consequences of mutating N851 in HIF-2ɑ only become apparent once T cells are allowed to differentiate *in vivo*, where they are guided by the varying microenvironments a T cell inhabits and transverses, including the tumor microenvironment. An in-depth analysis of such microenvironments would be necessary to determine how T cells expressing FIH-insensitive HIF-2ɑ gain a further advantage in rejecting tumors.

As with murine cells, human CD8+ T cells transduced with a CAR-HIF construct containing the PPA configuration showed increased cytolytic function against a B-cell lymphoma line. These results demonstrate that such vectors may be an important and novel addition to the CAR-T armamentarium.

## Acknowledgements

The authors thank Angelika Holler at University College London for providing reagents and discussions, and the Bioinformatics and Expression Analysis (BEA) Core Facility at the Department of Biosciences and Nutrition, which is supported by the board of research at the Karolinska Institute and the research committee at the Karolinska hospital.

The authors have declared that no conflict of interest exists.

## Supplementary Figures

**Supplementary Figure 1.**
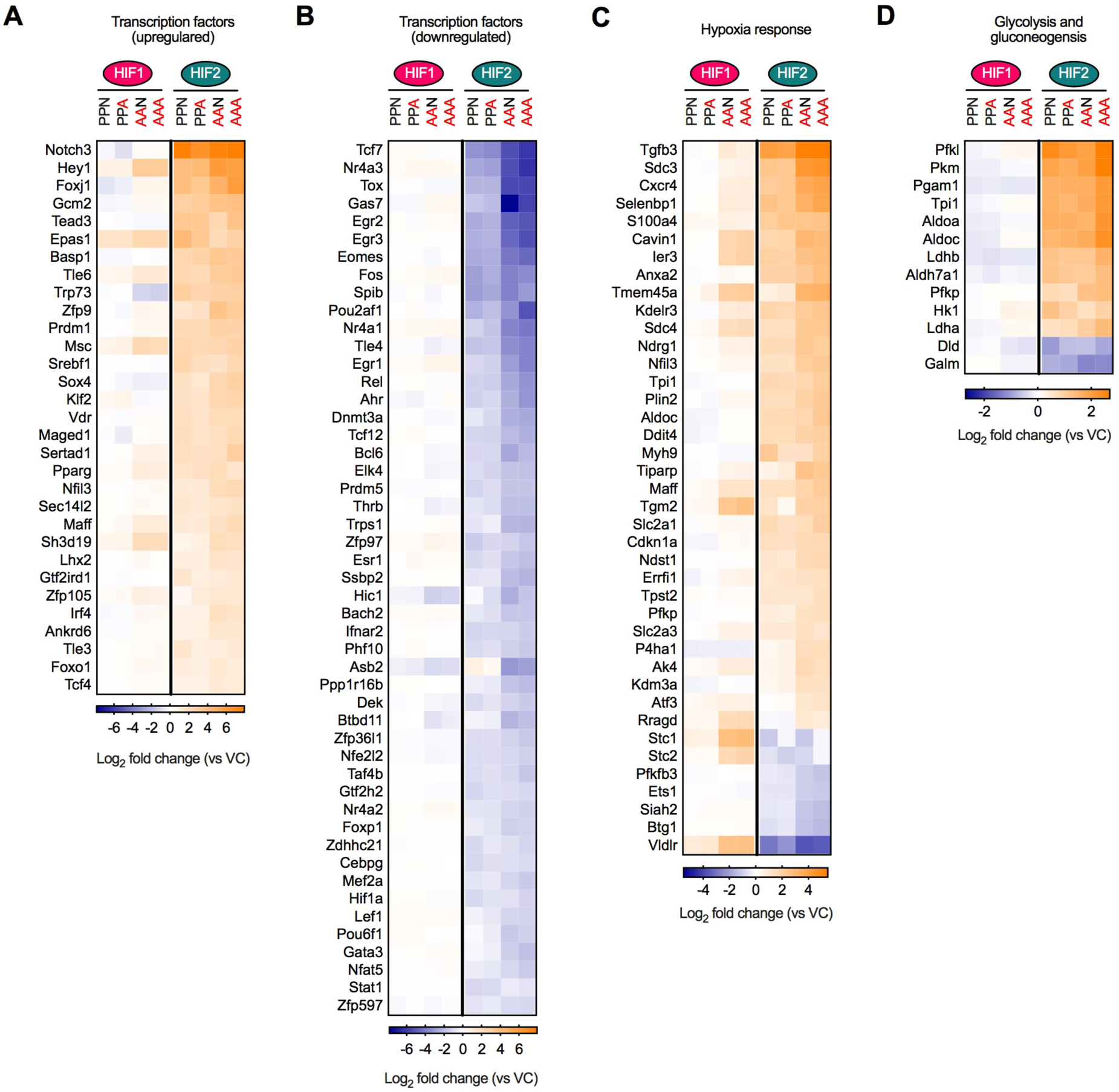
Ectopic expression HIF-2ɑ alters the transcriptional network of CD8+ T cells. **(A)** Heatmap showing Log_2_ fold change of upregulated transcripts involved in transcription modulation. **(B)** Heatmap showing Log_2_ fold change of downregulated transcripts involved in transcription modulation. **(C)** Heatmap showing Log_2_ fold change of deregulated transcripts involved in response to hypoxia. **(D)** Heatmap showing Log_2_ fold change of deregulated transcripts involved in glycolysis and gluconeogenesis. For each heatmap transcripts were included if FDR < 0.01 and Log_2_ fold change >1 or <−1 for at least one of the transduced groups.

**Supplementary Figure 2.**
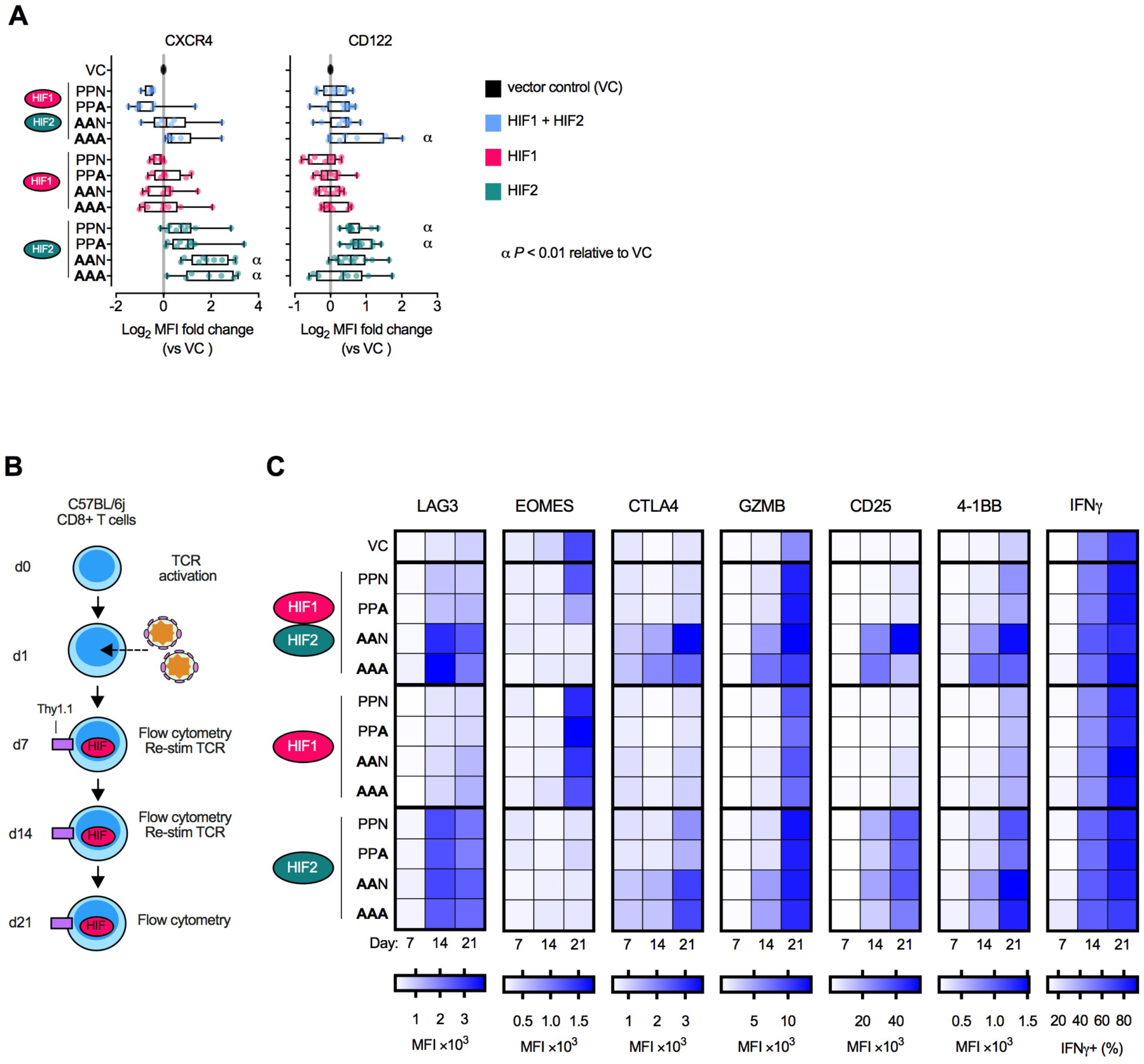
Ectopic HIF expression alters expression of proteins associated with CD8+ T-cell differentiation. **(A)** Expression of CXCR4 and CD122 determined by flow cytometry in CD8+ T cells transduced with vectors encoding HIF-1ɑ and HIF-2ɑ, HIF-1ɑ alone or HIF-2ɑ alone (day 3 to 5 post-transduction). PPN - inhibited by VHL and FIH; PPA - inhibited by VHL only; AAN - inhibited by FIH only; AAA - neither VHL nor FIH inhibition. Data expressed as Log_2_ fold change of median fluorescence intensity (MFI) relative to VC-transduced cells. Each data point represents an independent transduction (n=5-11). Results are pooled from a minimum of two independent experiments. α, *P* < 0.01; one-way ANOVA with Dunnett’s multiple comparison test relative to VC. **(B)** Time course of serially restimulated CD8+ T cells transduced with HIF-encoding vectors. Mouse CD8+ T cells were transduced with RV-vectors 24h after TCR activation. Phenotype of transduced cells was assessed by flow cytometry at days 7, 14 and 21 post-transduction. Transduced cells were restimulated at days 7 and 14 post-transduction. **(C)** Heatmap representing MFI or % IFNγ^+^ over time in transduced CD8+ T cells, as described in (B). IFNγ levels were measured after stimulation with PMA and ionomycin for 4 hours.

**Supplementary Figure 3.**
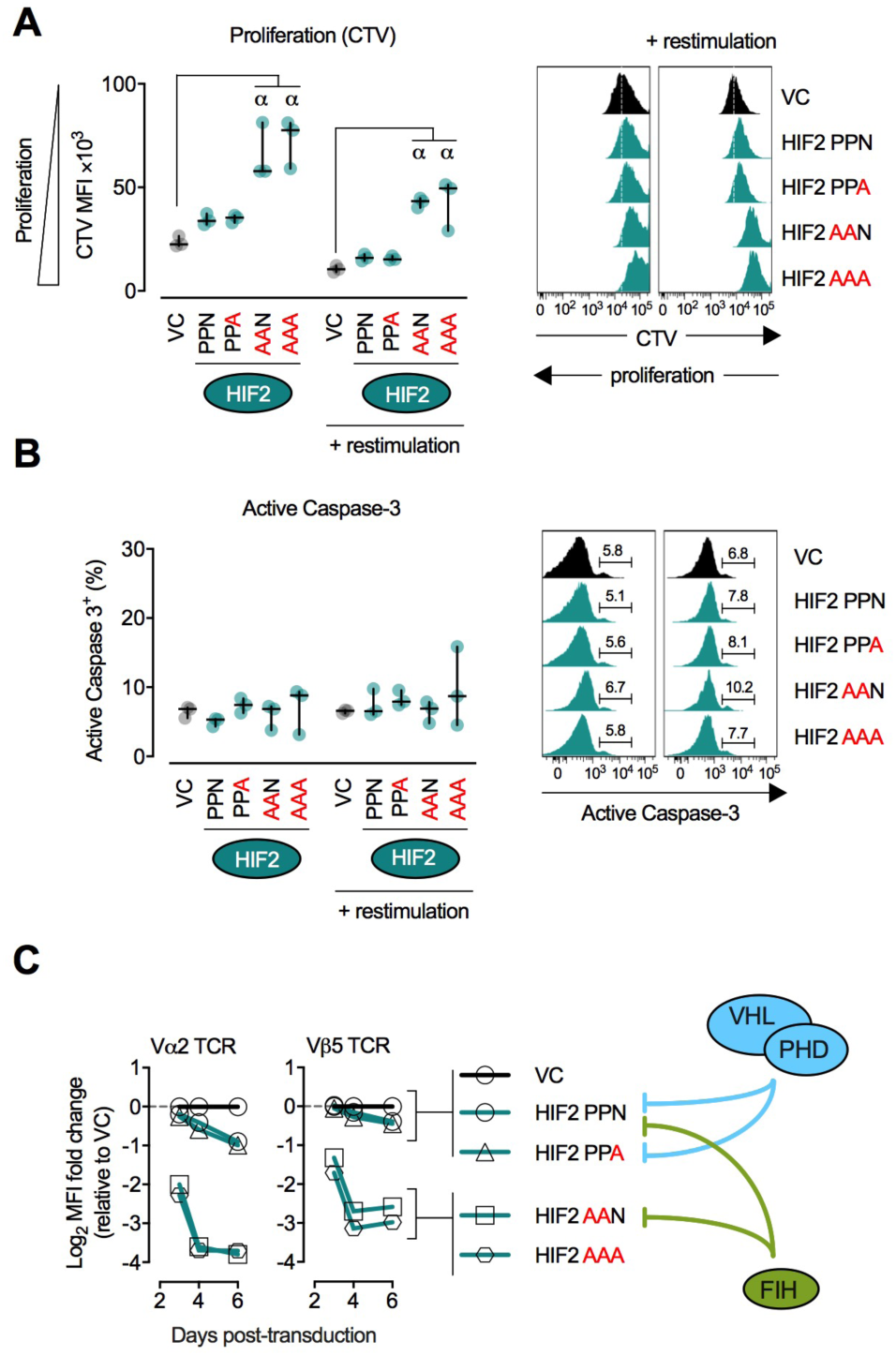
Ectopic expression of VHL-insensitive HIF-2ɑ reduces CD8+ T-cell proliferation and surface TCR expression while VHL-sensitive HIF-2ɑ increases expression of granzyme B. **(A)** CD8+ T cells were transduced with HIF-2ɑ-coding vectors and 6 days later were loaded with the proliferation dye cell trace violet (CTV) and either restimulated with αCD3/CD28 beads or left unstimulated for 3 days. CTV levels were determined by flow cytometry after gating on live, singlet, CD8+. Thy-1.1+ events. Left: summary data showing mean fluorescence intensity (MFI). n=3-4 independent transductions. Lines: median and interquartile range. Right: representative histograms. **(B)** As in (A) but for intracellular active Caspase-3 (marker for early apoptosis). **(C)** Progressive loss of surface TCR chains over time in HIF-2ɑ-transduced OT-I CD8+ T cells. Data are Log_2_ MFI fold change of TCR Vα2 and TCR Vβ5 chains up to day 6 post-transduction, relative to VC at each time point. α, *P* < 0.01; one-way ANOVA with Dunnett’s multiple comparison test relative to VC.

**Supplementary Figure 4.**
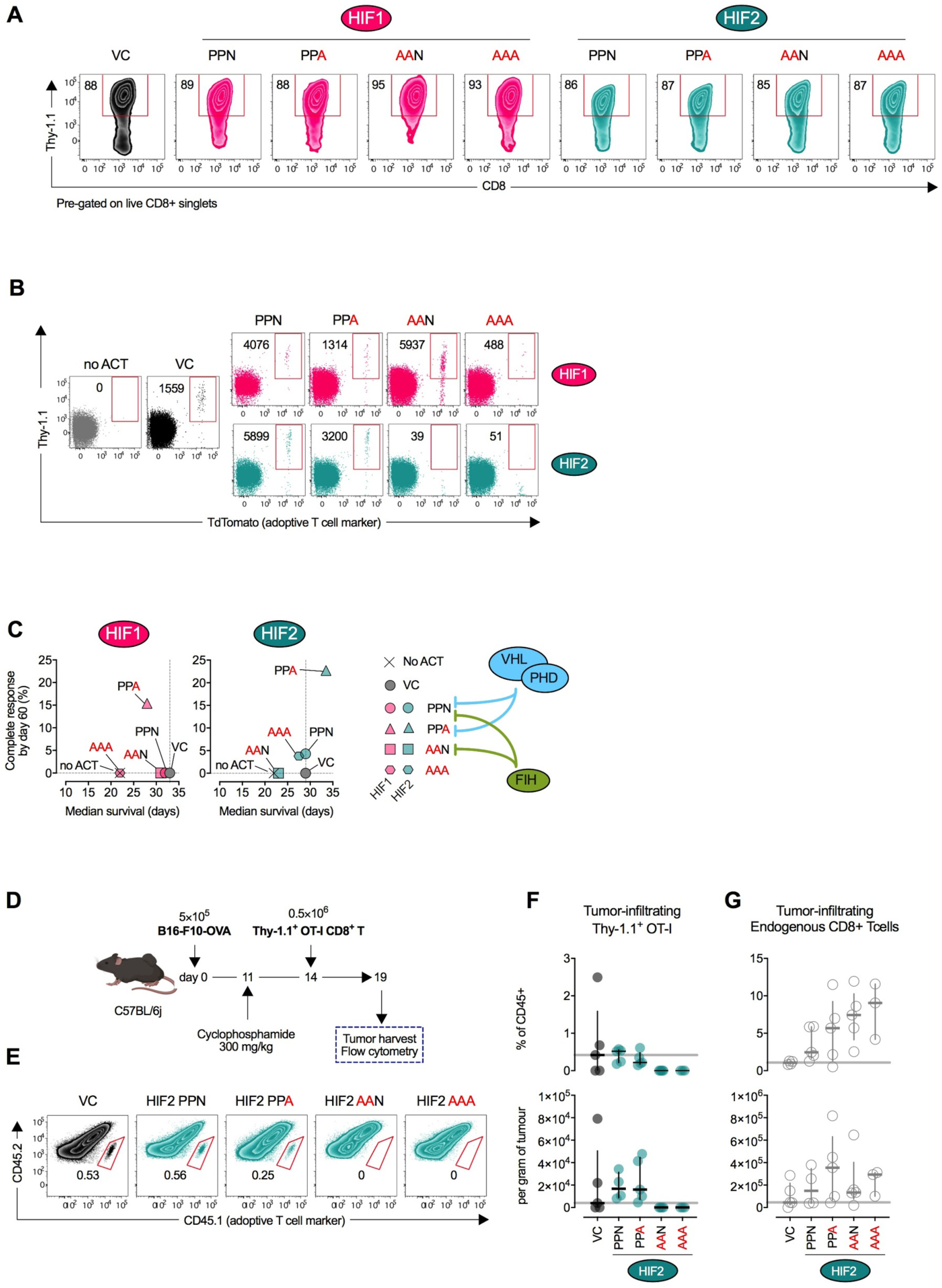
Adoptive cell transfer of HIF-1ɑ- and HIF-2ɑ-transduced OT-I cells into tumor-bearing mice. **(A)** CD8^+^ OT-I cells transduced with HIF-1ɑ- and HIF-2ɑ-coding vectors prior to adoptive cell transfer (ACT). 6 days after transduction cells were Thy-1.1 enriched using magnetic bead sorting and adoptively transferred shortly after into tumor-bearing mice. **(B)** Frequency of adoptively transferred HIF-transduced OT-I cells in peripheral blood of tumor-bearing mice at day 15. Representative flow cytometry dot plots (pre-gated on CD45+ live singlets) showing TdTomato (adoptive T-cell marker) and Thy-1.1 (transduction marker). Values are events per million PBMCs within the gate. VC: vector control. No ACT: no adoptive cell therapy. **(C)** Complete response rate (% of tumor-free animals at day 60) and median survival (extrapolated from survival curves shown in Figure 8E) for groups of animals receiving ACT of VC-, HIF-1ɑ- or HIF-2ɑ-transduced OT-I cells. Dashed lines: VC reference. **(D)** Adoptive cell therapy (ACT) model. C57BL/6j mice were injected subcutaneously with 5×10^5^ OVA-expressing B16-F10 (B16-F10-OVA) and 11 days later were lymphodepleted with 300 mg/kg cyclophosphamide. On day 14, 0.5×10^6^ HIF-2ɑ-transduced (Thy-1.1 enriched) OVA-specific OT-I CD8+ T cells were adoptively transferred. Tumors were harvested 5 days later, dissociated into a single-cell suspension and analysed by flow cytometry. **(E)** Frequency of adoptively transferred HIF-2ɑ-transduced OT-I cells within B16-F10-OVA tumors, 5 days after ACT. Representative flow cytometry dot plots (pre-gated on CD45+ live singlets) showing CD45.1 (adoptive T-cell marker) and CD45.2 (endogenous leukocyte marker). Values are the percentage of total CD45.2 events within the gate. **(F)** Frequency of tumor-infiltrating HIF-2ɑ-transduced OT-I cells as a percentage of CD45+ cells (top) or per gram of tumor (bottom). n=3-5 animals. Lines: median and interquartile range. Grey horizontal line: median for VC group. **(G)** As (F) but for tumor-infiltrating endogenous CD8+ T cells. α, *P* < 0.01; one-way ANOVA with Dunnett’s multiple comparison test relative to VC.

**Supplementary Figure 5.**
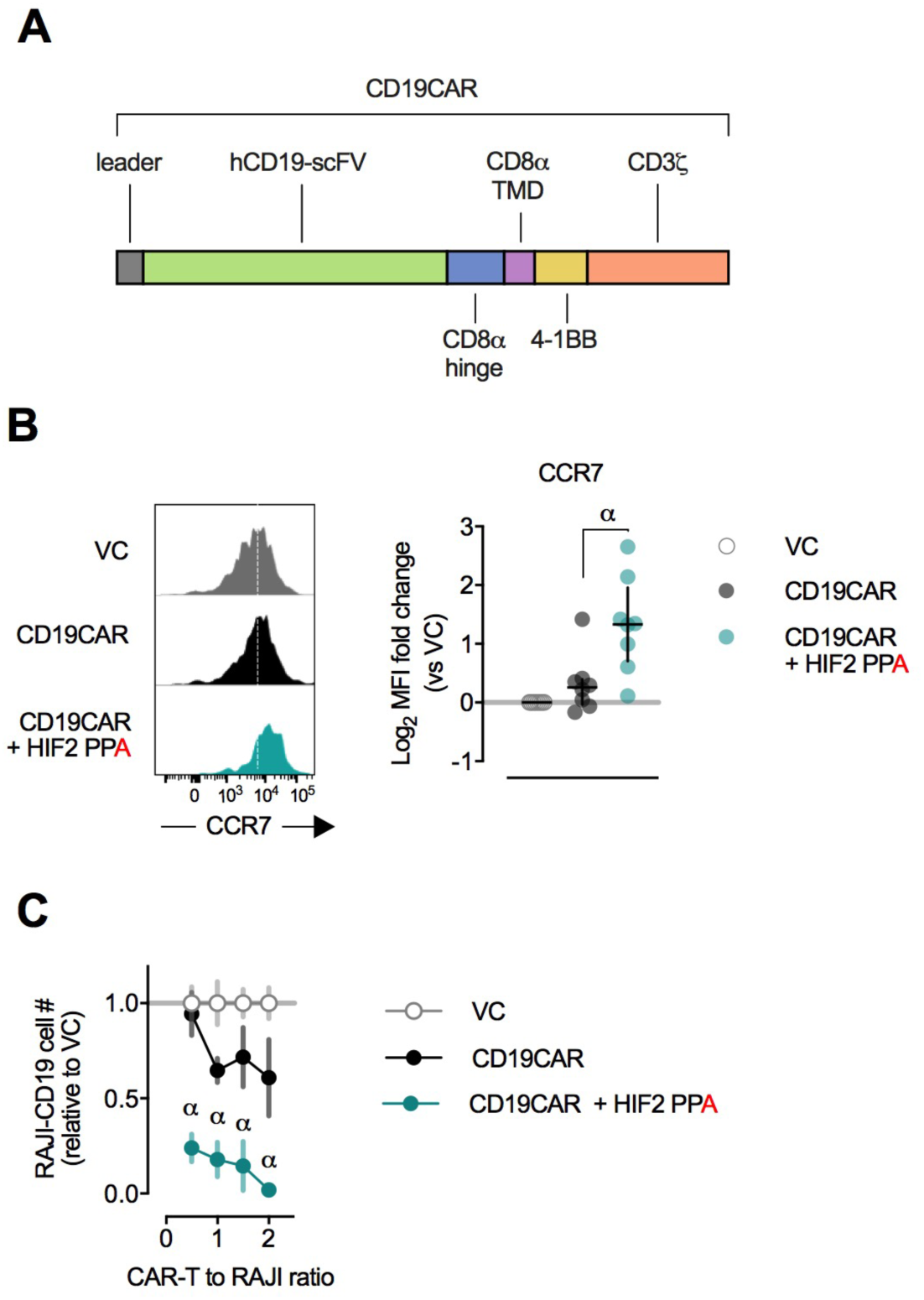
Ectopic expression of an anti-human CD19 CAR and FIH-insensitive HIF-2ɑ in human CD8+ T cells. **(A)** Structure of the anti-CD19 CAR used. scFV: single-chain variable fragment. TMD: transmembrane domain. 4-1BB and CD3ζ are the signaling domains from these proteins. **(B)** Surface expression of CCR7 in transduced CD8+ T cells, pre-gated on live, singlet, CD8+, GFP+ events. Representative flow cytometry histogram and summary data of Log_2_ mean fluorescence intensity (MFI) fold change versus vector control (VC). α, *P* < 0.01; one-way ANOVA with Dunnett’s multiple comparison test relative to VC. **(C)** Number of RAJI-CD19 after 20h co-culture with human CD8+ T cells transduced with VC, CD19CAR or CD19CAR and FIH-insensitive (PPA) HIF-2ɑ at varying CAR-T to RAJI ratios. n=5 human donors. RAJI counts are normalized to numbers in co-culture with VC T cells at each ratio. α, *P* < 0.01; two-way ANOVA with Dunnett’s multiple comparison test relative to VC.

## Protein Sequences

Peptide sequence for vector encoding mouse HIF-1ɑ and mouse HIF-2ɑ

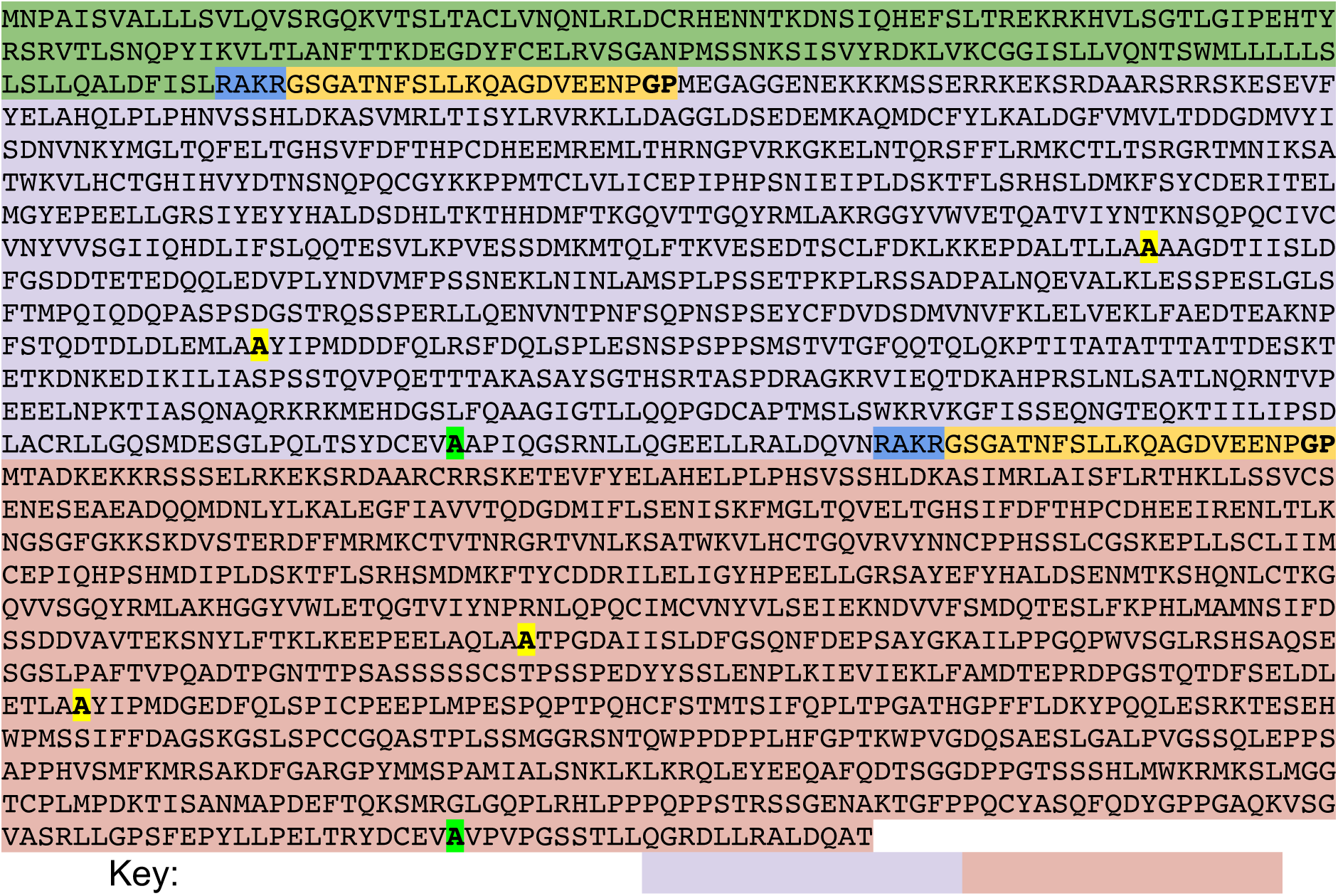

| Protein Name | NCBI Reference Sequence |
| --- | --- |
| Thy-1.1 | AAR17087.1 |
| Furin cleavage site | RAKR (cleavage after R) |
| Picornavirus 2A self-cleaving sequence | GSGATNFSLLKQAGDVEENPGP (cleavage occurs between G and P) |
| Mouse HIF-1α | NP_034561.2 |
| Mouse HIF-2α | NP_034267.3 |
| A Proline residue converted to Alanine |  |
| A Asparagine residue converted to Alanine |  |

Peptide sequence for vector encoding chicken ovalbumin (OVA)

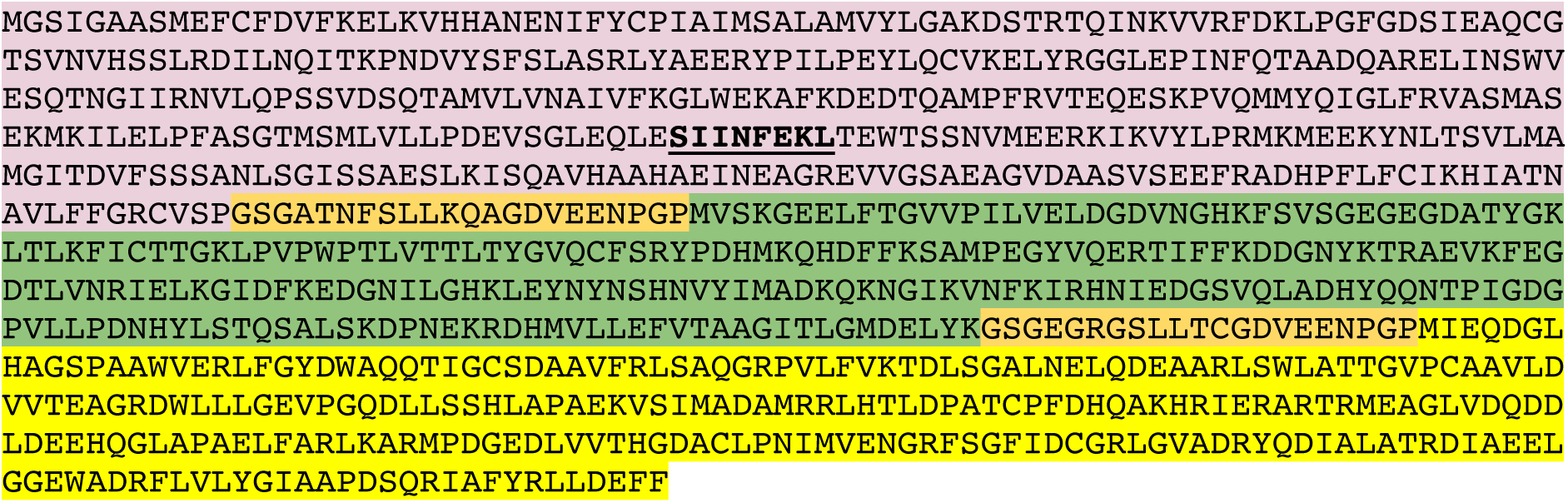

| Protein Name | NCBI Reference Sequence |
| --- | --- |
| Chicken ovalbumin | AUD54526.1 H2K-b-restricted OVA peptide (SIINFEKL) marked in bold. |
| Picornavirus 2A self-cleaving sequence | GSGATNFSLLKQAGDVEENPG**P** (cleavage occurs between **G** and **P**) |
| eGFP | ADQ43426.1 |
| Neomycin phosphotransferase | CAD21956.1 |

Peptide sequence for vector encoding CAR and human HIF-2ɑ

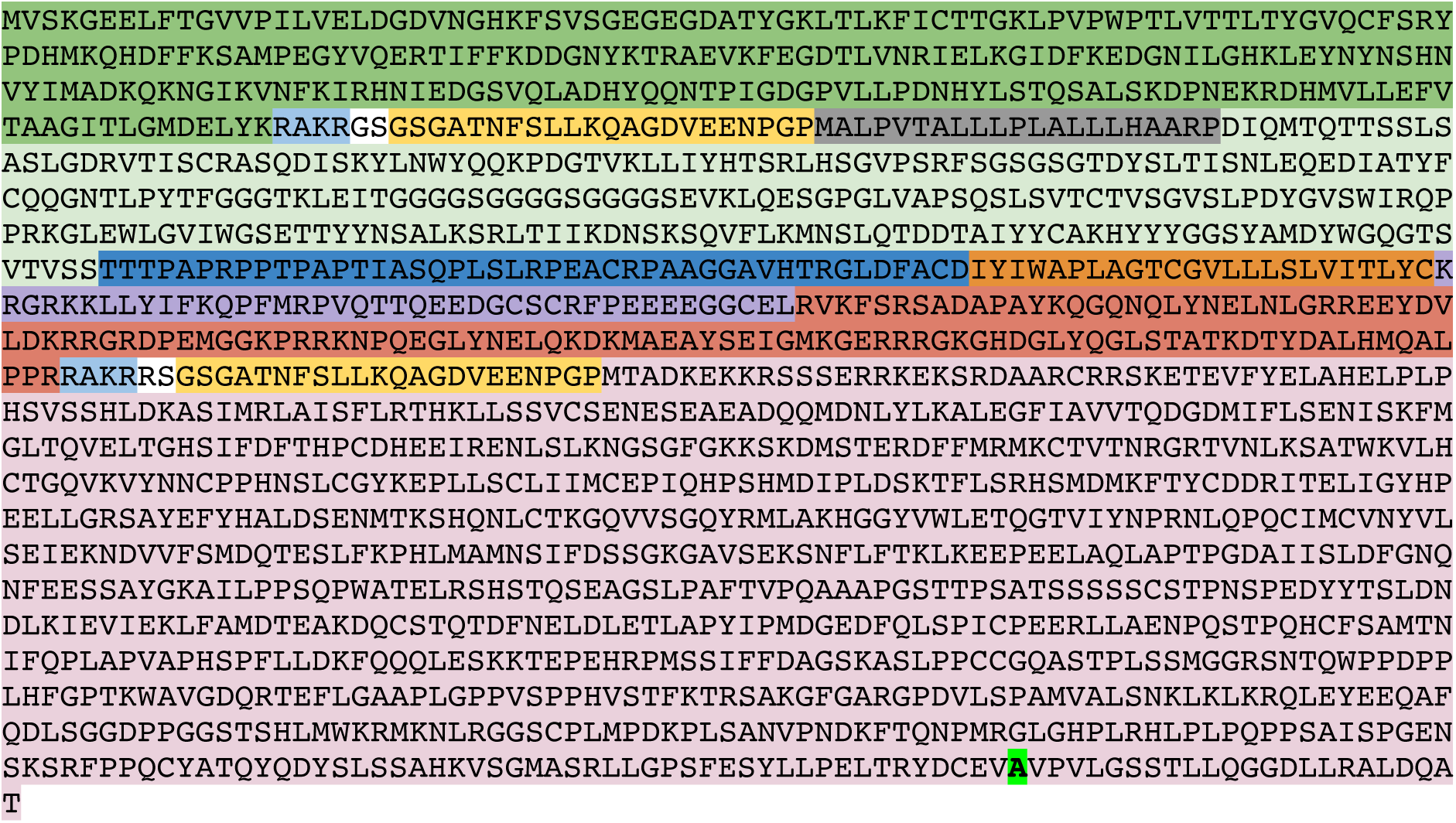

| Protein Name |  | NCBI Reference Sequence |
| --- | --- | --- |
| eGFP |  | ADQ43426.1 |
| Furin cleavage site |  | RAKR (cleavage after R) |
| Picornavirus 2A self-cleaving sequence |  | GSGATNFSLLKQAGDVEENPGP<br>(cleavage occurs between G and P) |
| hCD19 CAR | CD8a signal peptide | NP_001139345.1 |
|  | hCD19-scFv |  |
|  | CD8a hinge | NP_001139345.1 |
|  | CD8a TMD | NP_001139345.1 |
|  | 4-1BB | NP_001552.2 |
|  | CD3z | NP_000725.1 |
| Human HIF-2α |  | NP_001421.2 |
| A Asparagine residue converted to Alanine |  | N847A |

